# Memorization of novel patterns in working memory in a model based on dendritic bistability

**DOI:** 10.1101/2025.07.08.663796

**Authors:** Jiacheng Xu, Daniel L Cox, Steven J Luck

**Author notes:** Senior authors, listed alphabetically (Daniel L Cox, Steven J Luck).

## Abstract

Working memory can hold many types of information and is crucial for cognition. Commonly, models of working memory maintain information such as hues or words by forming memory attractors through structured connectivity. However, real-world information can be novel, making it infeasible to use predefined attractors. In addition, most models—with or without attractors—have focused on maintaining binary categories instead of continuous information in each neuron. In the present study, we investigate how the brain might maintain working memory representations of arbitrary novel patterns with graded values. We propose an unstructured, rate-based network model in which each neuron has multiple dendrites. Each dendrite shows bistable activity, which qualitatively captures the conductance-based dynamics in the corresponding spiking model and emulates fast Hebbian plasticity. This network can flexibly maintain novel graded patterns under various perturbations without fine-tuning of parameters. Through analytical characterization of network dynamics during the encoding and memory periods, we identify different conditions that yield either perfect memories or several types of memory errors. Our analysis reveals a functional separation for network neurons into two groups with distinct behaviors. Overall, this architecture provides robust and analytically tractable storage of novel graded patterns in working memory.

## Introduction

Working memory can actively maintain a variety of content types. Most previous studies have focused on simple content, such as location, color, shape, or letters [1, 2]. However, working memory can also hold complex content, such as a beach scene containing both basic elements (i.e., sand, cliff, sea or bird) and additional information (i.e., the inclined angle of the cliff or the speed of flying birds) [3]. The content can also be much more abstract, such as a new odor that elicits a distributed firing pattern in the cortex, or certain encrypted information during thinking. The underlying mechanisms of working memory for complex content are poorly understood.

By further elaborating the last example mentioned above, we formally define the theoretical question to study. Suppose someone is thinking about something, and he is interrupted by a temporary distractor, such as a loud noise, after which he resumes his thinking. During this short interruption, his thinking is paused, with relevant information stored somehow for quick retrieval when his thinking resumes. Although thinking itself involves complex neural dynamics, during the interruption, we assume that the paused information of thinking is stored actively in working memory, in a static, persistent pattern. As thinking can flexibly involve arbitrary information, this activity pattern is also arbitrary.

Therefore, in this paper, we explore the neural mechanism that allows the maintenance of arbitrary patterns, with each neuron having a specific firing rate across a range of possible values. Because there are exponentially many such patterns, the memorization should not rely on predefined attractors. This effectively requires a system to hold arbitrary novel patterns.

The majority of existing working memory models cannot memorize novel input patterns, because they require predefined attractors, such that only specific inputs limited to a predefined attractor space can be stored. For example, a line attractor can be constructed to store scalar inputs, like muscle strength, as activity intensities [4, 5, 6, 7], and a ring attractor can be constructed to store periodic input values, like color hues, as local neuronal activities along the attractor [8, 9, 10, 11]. All such working memory models need predefined attractors, with network connectivity explicitly structured or partially structured [11]. These attractors can be formed via extensive training, but they cannot maintain novel patterns unless they are similar to prior training examples. Conversely, maintaining novel patterns could potentially play a role in attractor formation. That is, holding novel patterns in working memory may provide the extended time needed for slow attractor formation and long-term storage [12].

There have been limited efforts to model the active maintenance of novel information. To perform this task, the system needs to: 1) have a certain form of fast plasticity, such that it can form an attractor in one shot based on a brief signal presentation, and 2) maintain stable memory activity in this attractor. Researchers have therefore proposed *fast Hebbian plasticity* [13], which is Hebbian plasticity but with faster changes in synaptic weight (within 100 ms). However, existing fast Hebbian plasticity models [14, 15] can only maintain binary memory activity, discarding any continuous information (e.g., input intensity). In addition, experimental evidence for fast Hebbian plasticity is sparse for working memory tasks [16, 17, 18]. However, the general concept of fast Hebbian plasticity does not have to occur in synaptic weights; it can instead be emulated by NMDA receptor-based dynamics. That is, researchers have suggested that these receptors effectively achieve fast Hebbian plasticity via the pre- and postsynaptic voltage dependence of NMDA receptor conductance. However, this activity-based, effective plasticity has only been used to maintain binary values and not patterns of graded activity [19, 20, 21].

In this paper, we propose a network model of working memory that can encode and maintain novel patterns with arbitrary graded firing rates across neurons. The model consists of a homogeneous network in which each neuron possesses multiple bistable dendrites. These bistable dendrites qualitatively emulate a form of fast Hebbian plasticity through dendritic activity. Each dendrite is either in an up-state or a down-state, but graded activation levels in the soma can be maintained by varying the number of bistable dendrites that are in the up-state. The same modeling idea can also be implemented for bistable synapses or bistable neurons, as described in the Discussion.

Although each modeled dendrite emulates fast Hebbian plasticity, our model contrasts sharply with classical models of long-term memory capacity. Our model is not about how many patterns can be passively stored in synaptic weights, but instead focuses on how to actively store a single novel pattern, with its content (arbitrary graded values) being more complex than that considered in most long-term memory models. Noteworthy, in classical long-term memory models [22, 23, 24], having only bistable values for synaptic plasticity is not desired for memory capacity, because this greatly reduces the scaling of memory capacity from linear, o(n_s_), to logarithm, o(log(n_s_)), with n_s_ represents the total number of independent synapses in a network. In contrast, in our model, bistable dendritic states in emulated fast Hebbian plasticity are crucial for obtaining an arbitrary graded memory pattern, as shown below. The focus of our model fits well with working memory, which is capable of holding complex information, but is limited to a small number of concurrent representations [25].

The remainder of the paper is organized as follows: First, we demonstrate the basic performance of the network and show its ability to robustly maintain information under various perturbations. Next, we characterize the detailed dynamics of the network during the encoding and memory periods. Then we analytically derive the input function for a perfect memory and identify mechanisms for various errors. These demonstrations use a simplified rate model, but we also demonstrate that similar results can be achieved by a more realistic spiking model. Finally, we discuss biological plausibility, different model interpretations, and comparisons with other models.

## Results

### Network structure and dynamics

We start by describing the rate-based network model. As shown in Fig. 1A, the network has N neurons that are all-to-all connected with weights equal to 1. Each neuron i has a firing rate f_i_ that obeys:

**Figure 1.**
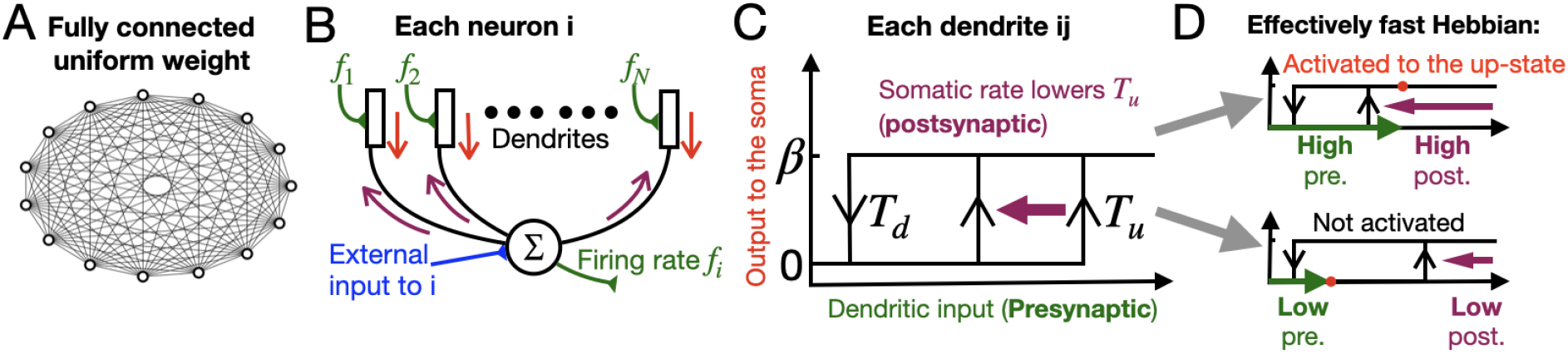
Network properties. **A**, Connectivity of the unstructured version of the network. N = 2500 neurons are fully connected with a uniform weight of w = 1. **B**, Design of simulated multi-compartment neurons used in the network. Each neuron has N separate dendritic units (or simply dendrites). Each dendrite receives an input from one other neuron in the network, and it also receives a backpropagated input from the soma. The dendrite’s output propagates to the soma. The firing rate of the neuron equals the summation of all dendritic outputs plus the external input to the soma. See Discussion for details about biological implementation and plausibility. **C**, Bistable dendrite input-output relation. Every dendrite exhibits the same bistability relation with an up-state value that occurs when the dendritic input exceeds an up-threshold T_u_ and a return to the down-state value when the dendritic input falls below a down-threshold T_d_. The up-state contributes a factor of β to the somatic firing rate. In addition, the somatic firing f_i_ can effectively lower T_u_. The effective up-threshold is modeled as 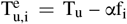, qualitatively capturing the effect of voltage backpropagation from the soma to dendrites. A spiking network implementation is provided later. **D**, Dendritic activation qualitatively emulates fast Hebbian plasticity. Left, when both pre- and postsynaptic firing rates are high, the dendritic input exceeds the low up-threshold, producing dendritic activation, with the stabilization point indicated by the red dot. Right, low pre- and postsynaptic firing rates cannot cause the dendritic input to exceed the up-threshold.

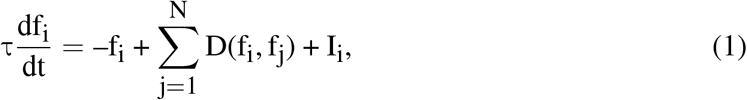

where the summation represents the total dendritic output, D(f_i_, f_j_) is the dendritic input-output relation, and I_i_ is the external input. As shown in Fig. 1B, each neuron has N identical dendrites, each of which receives a dendritic input from a separate network neuron j. Fig. 1C describes D(f_i_, f_j_): each dendrite can be either in an up-state or a down-state, with up- and down-thresholds T_u_ and T_d_. D(f_i_, f_j_) is mathematically expressed as

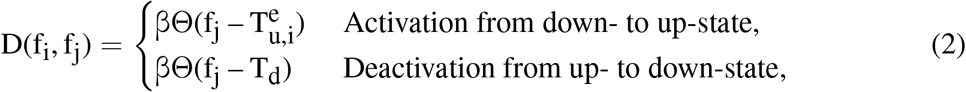

where β is the contribution factor for the up-state, Θ(x) is a Heaviside step function, and 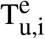 is the effective up-threshold. As illustrated by the purple arrow in Fig. 1C, the somatic firing f_i_ of a neuron can lower T_u_ to 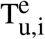 for all its dendrites, which we refer to as the *somatic effect*. This is modeled as

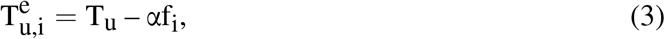

where α is the strength factor for the somatic effect. We further require 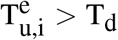.

The dendrite described in Eq. 2 resembles the dendritic performance of conductance-based models [19, 21]. The dendritic bistability can be produced by the dynamics of NMDA receptors. β qualitatively captures the increase in somatic firing due to high voltage transmitted from an activated dendrite (up-state). The somatic effect on the up-threshold is due to the back-propagated action potentials (bAPs) from the soma to all its dendrites. This effect is negligible for the down-threshold, so in the rate model, T_d_ is treated as a constant. We further constructed a spiking network with more details as mentioned below.

As shown in Fig. 1D, the activation of a dendrite from its down-state to its up-state is associative. That is, the activation not only depends on high dendritic input (presynaptic component), but also on a high somatic firing rate from the soma that the dendrite belongs to (postsynaptic component), which lowers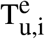. This makes dendritic bistability qualitatively resemble Hebbian plasticity with binary values. However, it is based on changes in dendritic activity rather than actual weight changes.

### Basic performance of the network

Novel graded working memory patterns can be maintained through bistable dendrites described in Eq. 2. During the encoding period, dendrites get activated. During the memory period, overall activity, and thus dendritic input, drops. However, if the dendritic input remains higher than T_d_, a dendritic up-state is still maintained. The firing rate at the soma depends on the number of dendrites in the up-state; an approximately continuous value in firing rate can therefore be maintained by maintaining a specific number of dendrites in the up-state.

Fig. 2A shows an example where a novel input pattern was memorized by this network. While the input pattern can be arbitrary, for visual clarity, we chose a pre-processed cloud image, with a graded set of values across 50 × 50 pixels (see further clarification at the end of this section). Each pixel maps to a separate neuron within the network (N = 2500 neurons). For the simulation, all dendrites were initially in the down-state. During the encoding period, the external input I_i_ received by each neuron was proportional to the corresponding pixel intensity. During the memory period after the external input was turned off, a stable activity pattern was maintained, with graded firing rates that formed a pattern similar to the original input.

**Figure 2.**
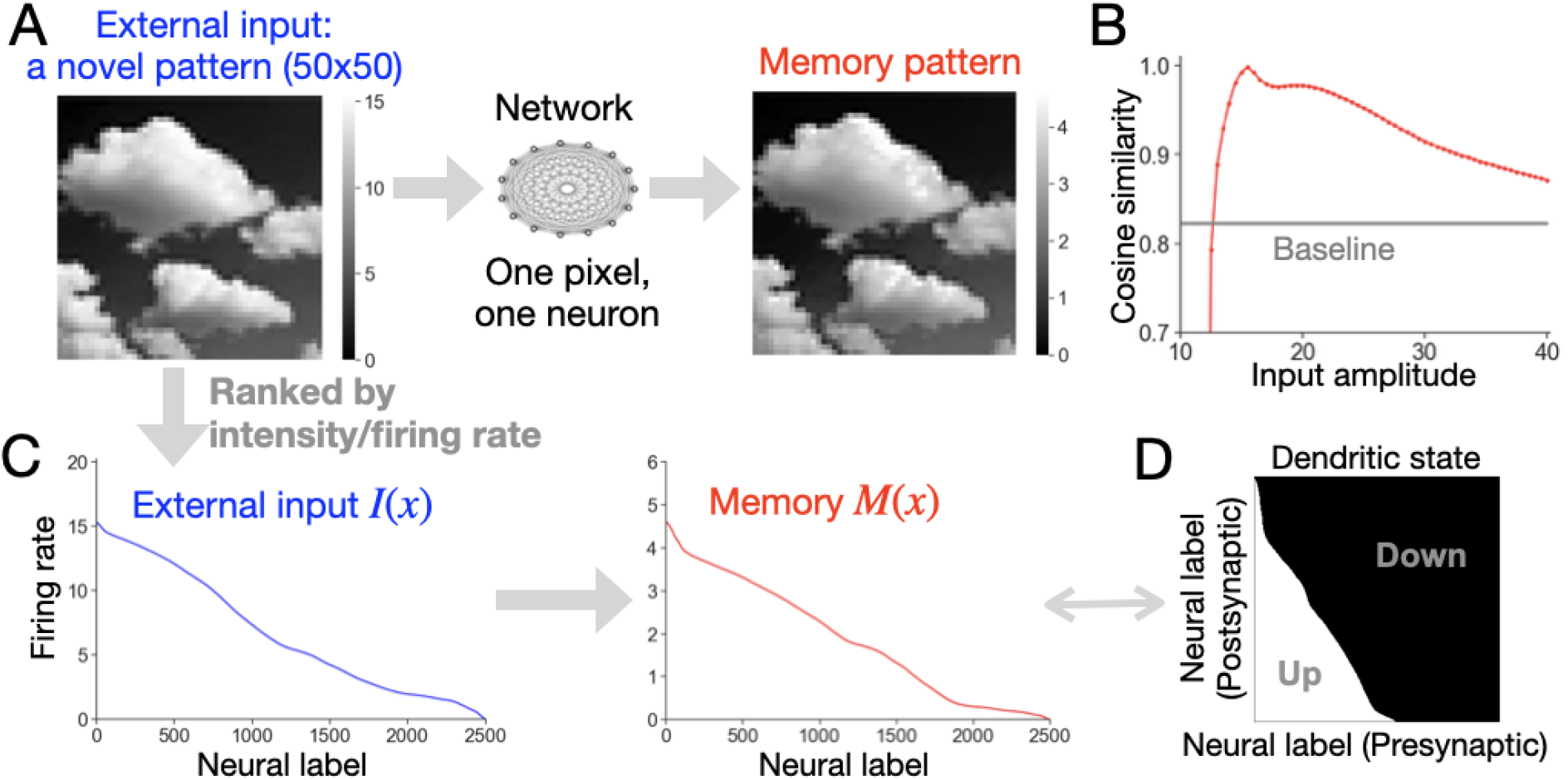
Memorization of an example novel pattern. **A**, Network performance for an input with a two-dimensional representation. Left, a graded cloud-like image (50 × 50) served as the input. Each pixel represents a neuron in the network, with the pixel’s intensity proportional to the neural firing rate. Right, when the input was removed, the resulting memory maintained a similar graded pattern despite some minor distortion. **B**, Memory performance across different input amplitudes. We changed the amplitude of the external input in *A* and recorded the corresponding memory pattern. The performance was evaluated by cosine similarity between the input and the memory. The baseline shows the performance if no graded information is stored in the memory (i.e., the cosine similarity between the input pattern and a flat, constant memory). **C**, One-dimensional representation of network activity, in which the neurons were ordered according to the input firing rate/intensity, in descending order. In this way, the external input and the memory can be described by two functions, I(x) and M(x), respectively. This reordering is justified because the neurons are functionally identical, and the original ordering was arbitrary. **D**, Dendritic states during the memory period. Each individual dendrite in the network is indicated by its pre- and postsynaptic neurons, which were ordered in the same way as in *C*. All dendritic states, with white for up and black for down, were plotted for the memory in *C*. The memory activity of a postsynaptic neuron is proportional to the total number of up-states across the corresponding horizontal line.

The network’s performance can be quantified by the cosine similarity between the input and the memory. Representing the two as two vectors 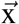 and 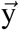, the cosine similarity measures the cosine value of the angle between the two, 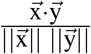. If input and memory match perfectly, up to a proportional constant, the value goes to one. As shown in Fig. 2B, we changed the amplitude of the overall input pattern, and checked the performance by cosine similarity. If the input amplitude was too small, only the high-firing portion of the input was memorized. If the input amplitude was too large, memory activity was saturated. The baseline (gray) shows performance if the memory is constant across neurons, which means that the network fails to memorize any graded information from the graded input. In this case, the cosine similarity is the same across input amplitudes for a fixed input pattern. See Appendix A for simulation details, and see below for a detailed analysis of the pattern of memory errors.

For ease of analysis, we can transform an input and the resulting memory into two one-dimensional vectors. For example, Fig. 2C shows two transformed vectors based on Fig. 2A. Because the present network is homogeneous, with each neuron anatomically equivalent, we can relabel the neurons in an arbitrary manner without loss of generality. It is convenient to relabel them in descending order according to the external input intensity, resulting in a monotonic vector of intensity values I_i_, which is approximated as a continuous function I(x). In addition, the network activity during the memory period is also a monotonic vector M(x) under the same set of ordered neural labels. This is because a larger input leads to a lower 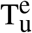, which activates more dendrites for memory. The detailed dynamics are explained below.

In this network, memory is not only represented in firing rates, but also equivalently in dendritic states. For the memory shown in Fig. 2C, Fig. 2D shows all the dendritic states with the same neural ordering. Each dendrite is represented by a point specified by the label of the neuron it belongs to (postsynaptic) and the neuron it receives a projection from (presynaptic), and each dendrite can be in the up-state (white) or in the down-state (black). Neurons that received greater pre- and postsynaptic inputs (i.e., those toward the lower-left side of the plot) have more up-state dendrites between them (i.e., more white points). This approach allows us to define *in-degree* as the number of up-state dendrites a neuron has, and *out-degree* as the number of up-state dendrites that the neuron projects to in other neurons. Note that in-degree value for a given neuron is proportional to its firing rate at equilibrium.

The simulations involved four additional key points. First, to approximate network performance in the continuum limit, we chose N = 2500. A novel pattern with 2500 entries may be too hard for humans, and 2500 separate dendrites in a neuron may not be possible. However, this large N makes it easier to illustrate the properties of the network. See the Discussion for alternative model implementations, which allow for more computational units. We also show a spiking model below with realistic settings. A second element of our simulations is that we used cloud images like the one shown in Fig. 2A as inputs; they are just examples of arbitrary novel graded patterns, and they were chosen simply for visual clarity. The analysis below is not related to any specific objects in an image. This model is about how to memorize a set of input values, not about representing a real visual image of a cloud, which would involve multilayer processing from the photoreceptors of the eye to the visual cortex. Third, the minimum external input was set to zero without loss of generality. Adding a constant I_c_ to the external input is equivalent to resetting the constant T_u_ to T_u_ – αI_c_. Finally, although the network can store graded input patterns, there can be an overall mismatch between the amplitude of the input pattern and the amplitude of the memory pattern, as shown in Fig. 2A,C. This can be captured by a multiplicative scalar factor, which may be implemented via a separate parametric working memory system [5, 26, 27]. In this work, we focus on the graded values of input patterns rather than this multiplicative factor.

### Memory robustness to perturbations

We next tested whether our model is robust to various perturbations without finely tuned model parameters. Perturbations considered include somatic noise, alternative connectivity patterns, and two-input interference.

We first considered robustness to small somatic Gaussian noise during the memory period. As shown in Fig. 3A, memory activity across neurons (dark red line) is, despite some fluctuation, nearly identical to the unperturbed memory (dotted red line). This robustness is based on dendritic bistability. The small noise changed the dendritic input, shifting the stable point (inset, red) back and forth, but this was insufficient to drive the stable point across T_u_ or T_d_. Consequently, the small noise did not cause dendrites to switch between up- and down-states, so the dendritic output to the soma remained unaffected.

**Figure 3.**
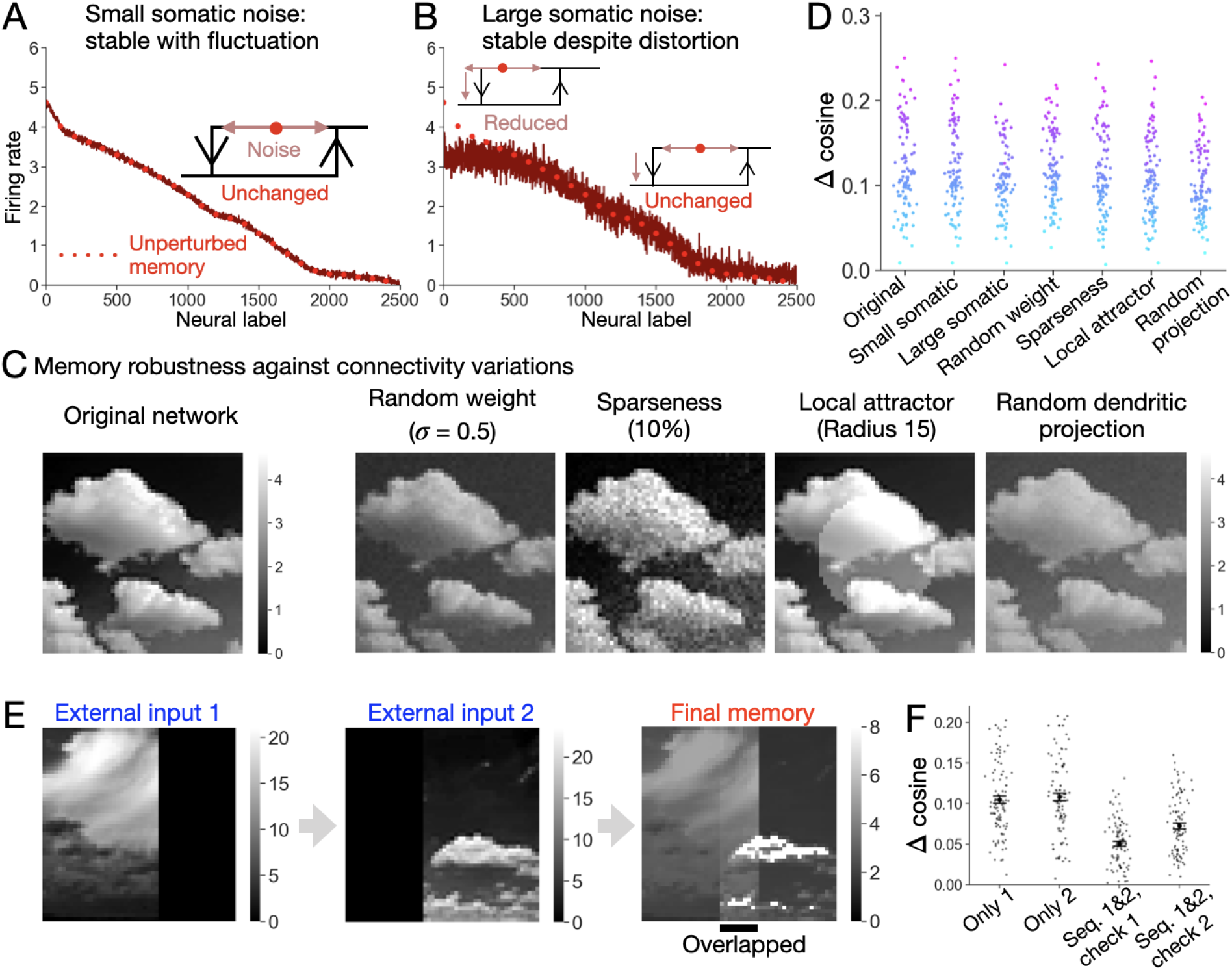
Memory performance under varying conditions. **A**, Network performance for the cloud image in the presence of somatic noise. Gaussian noise was applied to each soma independently. The dark red line shows the averaged memory over 500 ms. The dotted red line is the unperturbed memory replotted from Fig. 2 for comparison. Inset: A schematic plot of the bistability function illustrating how noise can drive a stable point in a bistable dendrite back and forth around its original stable point. **B**, Same as *A* but with larger noise. The noise caused the high-firing part of the memory to drop. Inset: Similar to *A*, but now some dendrites can flip to the down-state due to noise. **C**, Network performance under connectivity variations. Left: the unperturbed memory reproduced from the original network in Fig. 2. Right: memories from four connectivity manipulations: random weight values from a Gaussian distribution with mean one and σ 0.5; connection probability reduced from 100% to 10%; addition of a local attractor at the center; random projection for each neuron to dendrites of a targeted neuron, such that the same dendrite may receive multiple dendritic inputs. **D**, Statistical comparison across conditions for 100 simulated network inputs. The x-axis shows the conditions shown in *A-C*. The y-axis equals the cosine similarity minus the baseline value as shown in Fig. 2B. For each of the 100 input patterns simulated, we chose the input amplitude that maximizes the cosine similarity in the original network and applied this amplitude to all conditions. A given input pattern is represented by a consistent color across all conditions, determined by the cosine similarity observed in the ‘original’ condition. **E**, Interference between two overlapping input patterns. Two input patterns were delivered to the network sequentially. The first one was delivered to the left region, and, after an intermediate memory period, the second one was delivered to the right region, with the two overlapping in the middle. The final memory showed a graded pattern similar to both inputs. The middle overlapped region maintained graded values from both inputs, despite some interference. **F**, Statistical evaluation of the overlapping interference. 100 pairs of input patterns were simulated and recorded. The left two entries on the x-axis show the memory performance of the first/second input pattern, when only the first/second input pattern was delivered. The right two entries show the memory performance of the first/second input pattern, when two input patterns were delivered sequentially. Each error bar shows the respective standard error of the mean. See simulation details in Appendix A.

Fig. 3B shows an example of larger somatic noise. The memory (dark red line) still shows graded values but is more variable, with the high-firing neurons dropping their activities and relatively small changes to the low-firing neurons. From a dendritic perspective, negative noise fluctuation could cause some dendrites in high-firing neurons to transition from the up-state to the down-state, reducing firing rates (inset, upper). However, the same noise level did not change the dendritic states of low-firing neurons (inset, lower). Why did the activity reduction not lead to further recurrent activity reduction and eventually silencing the entire memory, via a positive feedback loop? Mechanistic details are explained in Appendix B, but the intuition is simple. As long as a recurrent activity reduction is not strong enough to drive a dendrite to deactivate by crossing, the dendritic output is unaffected.

Next, we examined memory robustness to variations in connectivity. For visual clarity, we chose the two-dimensional representation of the memories. Four conditions were considered (Fig. 3C, right). In the first case, the weights were changed from being exactly equal to one to being random values centered around one. In the second case, the network was sparsely connected instead of being fully connected. In the third case, a local attractor was added, implemented by using stronger connections between nearby neurons at the central region of the two-dimensional representation. In the fourth case, instead of requiring each neuron to project to a separate dendrite of the targeted neuron, projections were random, such that a dendrite could receive dendritic inputs from multiple neurons. In each case, the network robustly maintained a novel pattern similar to the original case. In particular, for the third case, the memory within the attractor still showed a graded pattern, indicating that this novel-input-storage mechanism can coexist with attractors in a single network. This property is consistent with the capability of the working memory system to hold both novel and familiar information.

Fig. 3D shows the statistical evaluations of these somatic noise and connectivity perturbations. 100 different cloud images were chosen as examples of arbitrary input patterns, each with an optimal input amplitude and a random initialization for noisy conditions. For each input pattern (colored), network performance was quantified as the cosine similarity minus the respective baseline. We found that the memories were typically quite robust to these perturbations. Only a small fraction of memories (about 10 across all perturbations) gave negative performance, meaning worse than a uniform memory.

The interference between two sequentially encoded memories was examined in Fig. 3E,F. After input 1 was memorized in the network, another partially overlapping input 2 was delivered, making the final memory similar to both inputs, as shown in Fig. 3E. Errors can arise in two ways. First, the high-firing neurons for the existing memory from input 1 can erroneously activate dendrites during the encoding period of input 2, regardless of overlap. Second, for the overlapped region, the final memory showed an additive pattern based on both inputs. This pattern mixture in the middle could be hard to separate, such that we might not know whether a memory value belongs to input 1 or 2. In Fig. 3F, we ran 100 pairs of input patterns and recorded the performance. For comparison, the performance of input 1 (or 2) when it was the single input was also recorded. Overall, memory shows robustness to the two-input interference. For the purpose of this paper, we focus on the working memory flexibility of a single pattern, not on capacity, so we did not explore multiple working memory patterns further. See Appendix A for more simulation details.

### Detailed network dynamics

How are graded patterns actually memorized by this network? In this section, we provide a detailed breakdown of the network dynamics during the encoding and memory periods. For the dynamics during the encoding period, Fig. 4A-C is designed to provide intuition. Neurons in Fig. 4A are ranked in descending order as in Fig. 2C. For a particular neuron A (Fig. 4B), all its dendrites receive graded dendritic inputs recurrently and share the same effective up-threshold 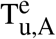, which is lower for larger somatic firing f_A_. A dendrite is activated if its dendritic input exceeds 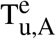. Counting all activated dendrites gives the summed dendritic output, up to an up-state contribution factor β (Fig. 4C). This is added to the soma along with the external input to get a new firing rate, which is iteratively fed back in Fig. 4A. Overall, each neuron receives identical but graded dendritic inputs. What differentiates the activation in each neuron is the somatic effect. For example, a neuron with a higher firing rate has a lower 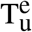 for all its dendrites, allowing more dendrites to activate. This eventually leads to a stable but graded firing rate.

**Figure 4.**
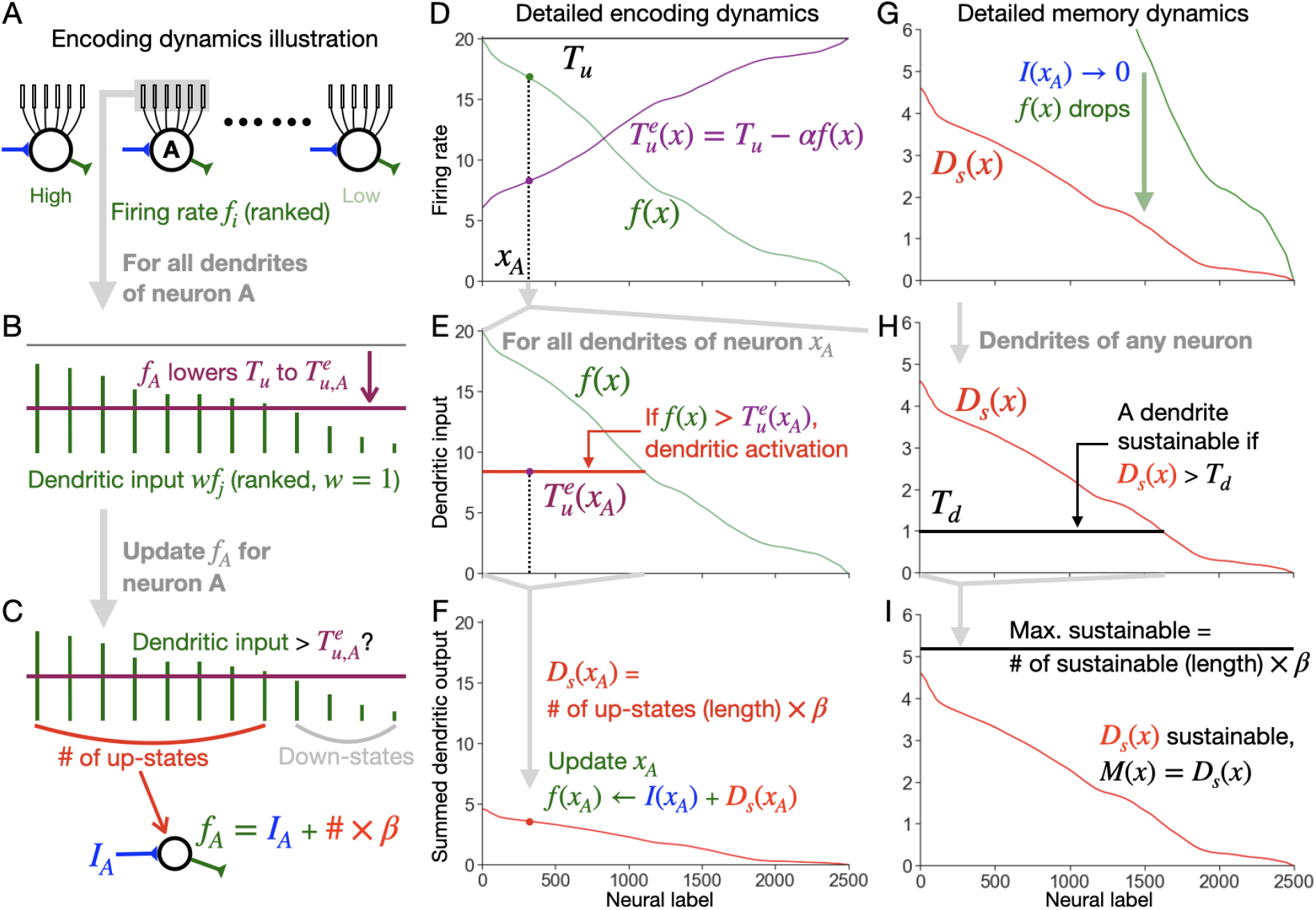
Detailed dynamics during the encoding and memory periods. **A-C**, Illustrative plots of the encoding dynamics. **A**, Network neurons ranked by external input intensity. **B**, All dendrites of neuron A. Dendrites are ordered by the input intensity of their presynaptic neurons. Dendritic inputs with graded values (green sticks) go through a weight of w = 1. The somatic effect uniformly lowers T_u_ to 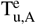 (purple bar) for all dendrites of neuron A. **C**, Dendritic activation. If a dendritic input exceeds 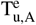, the dendrite is activated. Each activation contributes β to the somatic firing. All dendritic contributions are summed in the soma along with the external input (blue). Down-state dendrites have no effect. **D-F**, Detailed encoding dynamics, drawn to illustrate Eq. 4. The illustrative curves here are based on the stabilized activity in Fig. 2C, while the dynamics shown are generally true. Similarly below. **D**, The firing rate f(x) (green) effectively lowers T_u_ to the effective up-threshold 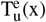 (purple). **E**, For any neuron x_A_, all its dendritic inputs, with weight w = 1, are plotted. All dendrites in x_A_ have the same 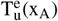. The portion of dendrites activated is shown by the red line. **F**, Value updating for each x_A_. The summed dendritic output D_s_(x_A_) equals the number of activated dendrites times β. This then updates the firing rate: f(x_A_) = D_s_(x_A_) + I(x_A_), with the external input I(x_A_). f(x_A_) is iteratively fed back to *D* until the system is stabilized. **G-I**, Detailed memory dynamics, drawn to illustrate Eq. 5. **G**, With I(x) turned off, D_s_(x) remains. Note that the y-axis is rescaled from *F*. **H**, Unlike 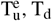 is a constant. The black line shows the portion of dendrites with dendritic inputs above T_d_. **I**, The maximally sustainable activity (black line) is given by the number of sustainable dendrites times β. If D_s_(x) falls below the line, as is the case here, D_s_(x) is sustainable, equaling the memory M(x). See details in the text.

Next, we formally describe the encoding dynamics in the continuous limit. During the encoding period, firing rates increase monotonically in time, such that the dendritic bistability relation reduces to a step function with a threshold T_u_. The firing rate f(x) (green) and the corresponding 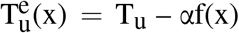 (purple) are shown in Fig. 4D. For a neuron x_A_, Fig. 4E shows all its dendritic inputs wf(x), with w = 1. If a dendritic input exceeds the effective upthreshold, 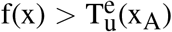, the dendrite is activated. As f(x) is monotonic, the number of up-state dendrites equals the length of the red line, 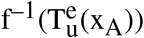, where we assume the existence of the inverse function f^−1^(x). Next, as shown in Fig. 4F, for each label x_A_, the summed dendritic output D_s_(A) is updated as 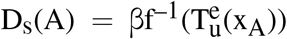. This thus gives an updated firing rate, f(x) = I(x) + D_s_(x), with I(x) denoting the external input. f(x) is iteratively fed back into Fig. 4D until the system is stabilized, resulting in an overall dynamical equation:

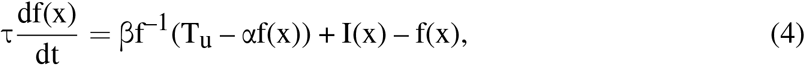

with a time constant τ to capture the slowest dynamic timescale. We denote the stabilized firing rate during the encoding period as F(x).

Following that, we describe the dynamics during the memory period. With I(x) turned off in Fig. 4G, f(x) reduces to D_s_(x). But can D_s_(x) self-sustain as memory activity or will it also decrease, leading to forgetting? To answer this question, in Fig. 4H, we zoom in on all dendrites of a neuron x_A_. An activated dendrite can still be sustained if its dendritic input exceeds T_d_, i.e., D_s_(x) > T_d_. If all activated dendrites satisfy this condition, then D_s_(x) is stable without forgetting. As D_s_(x) is monotonic, the maximal number of sustainable dendrites equals the length of the black line, 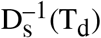. Consequently, the maximal sustainable activity is given by 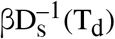, as shown by the black line in Fig. 4I. If 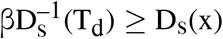, all activated dendrites are sustainable, and the memory is M(x) = D_s_(x). If 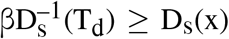 is not satisfied, memory forgetting occurs. Some once-activated dendrites are no longer sustainable, due to insufficient dendritic inputs, and fall back to the down-state. This process follows the equation:

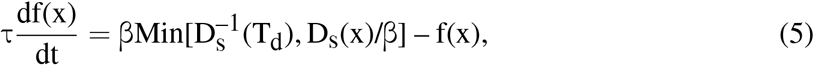

where the memory M(x) is given by the stabilized f(x) during the memory period. We primarily focus on the performance when there is no forgetting. See Appendix B for a detailed examination about memory forgetting.

### Perfect memory and the functional separation of two groups

With the dynamics in the Eqs. 4 and 5, the network can maintain a perfect memory. Although biological working memory systems are not typically capable of perfect memory, the present analysis can gain some mechanistic insights.

By perfect memory, we mean that a memory M_p_(x) is proportional to the input I_p_(x) by a multiplicative factor. Denoting the stabilized firing rate during the encoding period as F_p_(x) and the summed dendritic output at this time as D_s,p_(x), we obtain M_p_(x) = D_s,p_(x) *∝* I_p_(x) *∝* F_p_(x).

During the encoding period, the proportionality requires the stable state to satisfy:

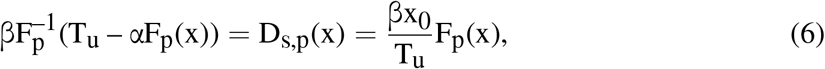

where 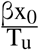 is the proportionality ratio, with an arbitrary neural label x_0_ between 0 and N. The reason for this particular form is explained below. During the memory period, all activated dendrites should be sustainable for memory.

Based on the detailed algebra in Appendix C, a perfect memory needs the input to satisfy:

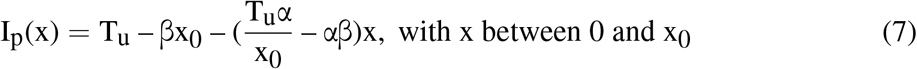

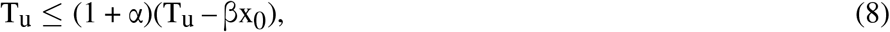

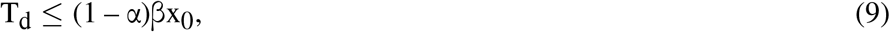

where the non-negativity of I_p_(x) requires 1 ≥ α > 0. The first inequality ensures that the input is large enough to activate dendrites, and the second inequality ensures that no forgetting happens during the memory period. This solution shows that I_p_(x) needs to be partially tuned: for x between 0 and x_0_, I_p_(x) is linear; for x between x_0_ and N, I_p_(x) can be arbitrary. A particular I_p_(x) is shown in Fig. 5A. The location of x_0_ is marked by the cyan dot. M_p_(x) perfectly matches I_p_(x) after rescaling. This holds true for arbitrary I_p_(x) values within the gray box. See Appendix A for simulation details.

**Figure 5.**
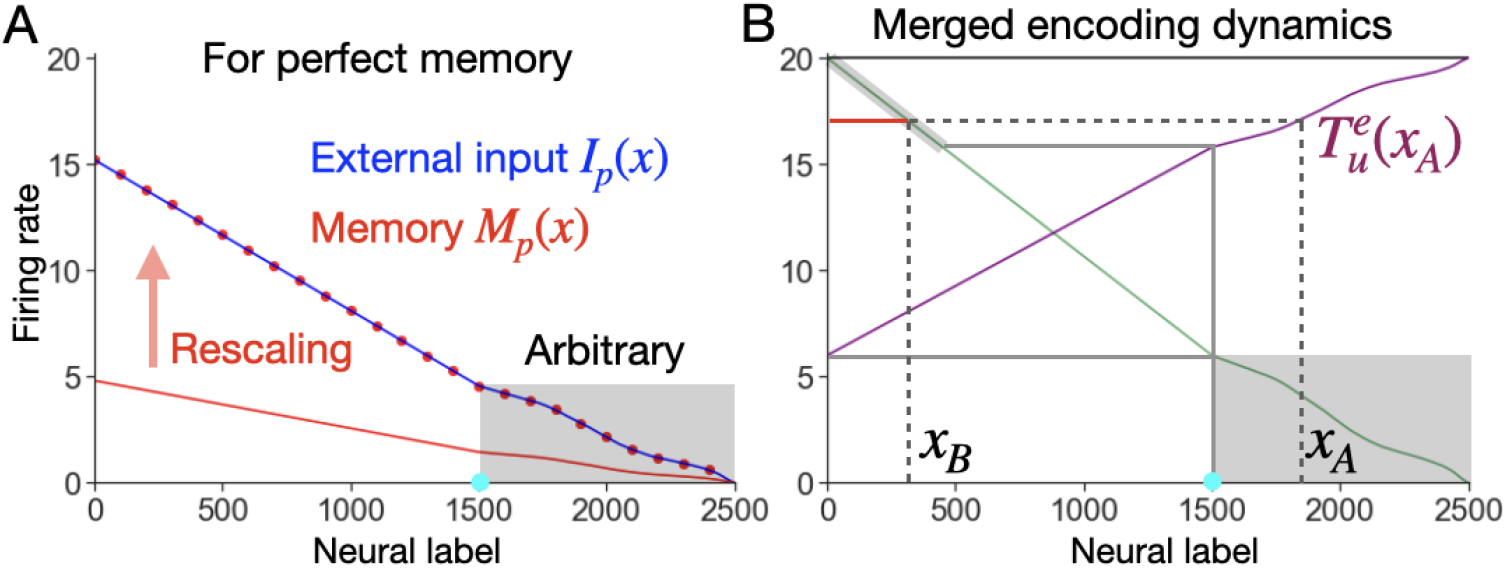
Perfect memory (up to a multiplicative factor). **A**, Network performance. A particular form of external input I_p_(x) (blue) gives a perfect memory M_p_(x) (red line) up to a multiplicative factor. M_p_(x) perfectly agrees with I_p_(x) after rescaling (dotted red line). Given x_0_ (cyan dot), I_p_(x) needs to be linear for neurons on the left side of it, while I_p_(x) can be arbitrary on the right side of it (gray box). **B**, Dynamics during the encoding period. This plot merges several steps, as illustrated in Fig. 4D-F, into a single one for simplicity. The cyan dot in *A* is graphically determined. See details in the text.

Eq. 7 reveals two functionally separate groups of neurons. The high-firing group requires a linear relationship with firing rates, while the other low-firing group is unconstrained. The formation of two groups during the encoding period is explained in Fig. 5B, where we merged all three dynamic steps, as shown in Fig. 4D-F, into a single one. Neurons with high firing rates can activate the dendrites they project to (out-degree). Such activation can lead to further interaction. Therefore, for a perfect memory, firing rates need to be well-tuned to prevent interaction. See the next section for why firing rates need to be linear. In contrast, neurons with rates below the smallest 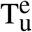 at x = 0 cannot activate any dendrites they project to (no out-degree). Because of this, they do not affect other neurons, such that they can receive unconstrained inputs. Formally, we define these two functional groups as the *noninteractive group*, with no out-degree, and the *interactive group*, with out-degrees. They are separated by the cyan dot, which is determined by the extended line from the smallest 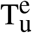 value as shown graphically.

These two functional neuron groups generally exist beyond the perfect case. For example, in Fig. 2D, the value of out-degree decreases from left to right along the horizontal axis, until it hits zero. This separates neurons into the interactive group (left) and the noninteractive group (right). These two functional groups also behave differently when error occurs, as discussed below.

### Different errors from the encoding period

If I(x) does not follow the exact form in Eq. 7, a graded memory can still be formed, but with errors. In this section, we examine several types of errors that originate during the encoding period. For errors due to forgetting during the memory period, see Appendix B.

### In-degree error, out-degree error, and multiselectivity

Ideally, if there is a local increase in the external input one neuron receives during the encoding period, this should cause an increase in the memory activity of that neuron, with a magnitude proportional to the input increase. If this is true and is applied to each neuron, any graded input can be memorized perfectly, up to a proportionality constant. However, for this network, the memory increase is not just determined by the local input increase (*in-degree error*), and the memory response may exist beyond the location where a local input is applied (*out-degree error*). Below, we first show the example and the mechanism for each error, then discuss the interaction between the two errors.

Let’s look at in-degree error first. Fig. 6A shows an example in which a local input increase leads to an extra memory response. However, the magnitude of this response was not proportional to the magnitude of the input increase. The mechanism is explained in Fig. 6B with the merged encoding dynamics. Here, we treat the input increase as infinitesimal, so that we can ignore its width and strength. Based on the dynamics shown in Fig. 4D-F, in Fig. 6B, the number of activated dendrites (in-degree) for neuron x_A_ is proportional to the length of the red line. An input increase to x_A_ lowers its up-threshold 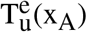, increases its in-degree, and thus increases the summed dendritic output D_s_(x_A_) and its memory M(x_A_). This increase depends on f^*′*^(x_B_), the slope of f(x) at x_B_. This property is unwanted, as this makes the memory response for x_A_ determined by a nonlocal slope at x_B_, causing the in-degree error. This error exists in all neurons having nonzero memory activity (nonzero in-degree).

**Figure 6.**
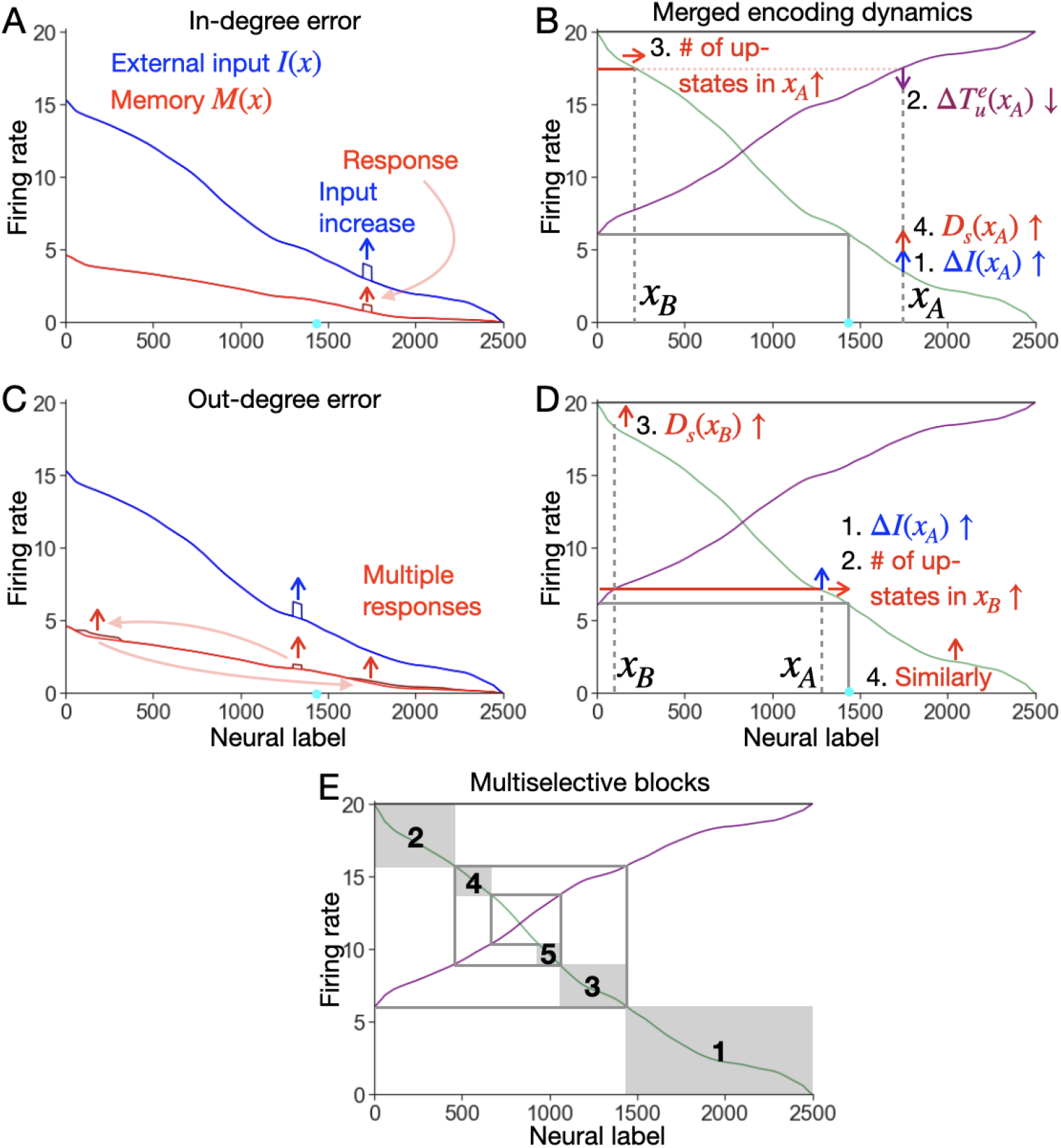
In-degree and out-degree errors in memory. **A**, In-degree error. With an extra input increase delivered locally to some neurons during the encoding period, the memory response due to this extra signal is local, but with a magnitude not proportional to the strength of the extra input. **B**, Merged encoding dynamics of the example in *A*. This demonstrates the mechanism underlying the in-degree error. **C**, Out-degree error. With an extra input increase delivered locally to neurons in the interactive group (left-hand side of the cyan dot) during the encoding period, the memory responses show up in three locations (red arrows). **D**, Merged encoding dynamics of the example in *C*. This demonstrates the mechanism underlying the out-degree error. **E**, How extra input leads to interaction during the encoding period. Neurons are functionally separated into different regions, with region 1 being just the noninteractive group. These groups are determined by the spiraling gray line starting from the point of the smallest 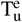 value. See detailed explanations in the text.

In contrast, out-degree error only exists in the interactive group, where neurons have nonzero out-degrees. An example is shown in Fig. 6C, where a local input increase to neurons in the interactive group caused memory changes in three locations. The mechanism is explained in Fig. 6D. A neuron x_B_ receives dendritic inputs, some above 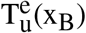 and some below. There exists a dendrite whose dendritic input is right below 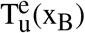. We denote the presynaptic neuron of this dendrite as x_A_. An increase in f(x_A_) can activate the corresponding dendrite in x_B_ (in the out-degree of x_A_), inducing a memory increase nonlocally in x_B_. Similarly, the increase in x_B_ can trigger a nonlocal memory increase in another neuron by the same mechanism (red arrow in the lower right corner in Fig. 6D). This mechanism can sequentially induce nonlocal memory increases until it stops at a neuron in the noninteractive group, where no out-degree exists. In addition, in parallel with the nonlocal out-degree error, the magnitude of the memory change in each neuron is subject to the in-degree error, as explained above. In Fig. 6C, because the local input increase was not infinitesimal but had finite width, the memory responses also varied in width.

In-degree and out-degree errors give a particular form of multiselectivity. In Fig. 6E, the firing rate during the encoding dynamics can be separated into different regions (gray boxes) as determined by the spiraling gray line starting from the lowest 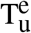. Each gray box includes some neurons (along the x-axis) depending on their firing rates. An infinitesimal input increase in region 3, as is the case in Fig. 6D, gives three memory responses, located in regions 3, 2, and 1. More generally, a neuron in a region R can influence in a total of R regions, R, R – 1, …, 1, while it can also be susceptible to change to an input increase in regions R + 1, R + 2, …. That is to say, for a neuron with a smaller region label R, it is less influential (inducing less nonlocal changes to other neurons) but more susceptible to a small extra input increase (e.g. from noise) from neurons in other regions. And neurons with a larger region label R are more influential but less susceptible. Overall, neurons exhibit a level of anticorrelation in their influence and susceptibility levels, and the specific influence and susceptibility levels depend on the region label each neuron belongs to, which is further determined by the firing rate of each neuron. Such a special form of multiselectivity among neurons can be potentially tested in the brain by perturbing neural activities.

With in-degree and out-degree errors introduced, we revisit the perfect memory case in Fig. 5 for a deeper understanding. Recall that a perfect memory requires the stabilized firing rate during the encoding period F_p_(x) to be linear in the interactive group and arbitrary in the noninteractive group, as shown in Eq. 7. Why does this particular form of input lead to a proportional memory activity? For any neuron x_A_ in the noninteractive group (Fig. 5B, shaded box, R = 1 using the label in Fig. 6E), its memory change is nonlocally proportional to 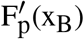, within the thick gray line (R = 2). To ensure that any input to x_A_ induces a proportional memory change, we need to remove this nonlocal dependency of the memory in x_A_ on 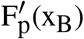. This requires 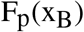 to be a constant, or F_p_(x_B_) to be linear. Similarly, to ensure any input to x_B_ induces a proportional memory change, we need to remove the nonlocal dependency of the memory in x_B_ on 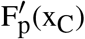, with x_C_ existing in region R = 3 by extending the spiral line in Fig. 5B as was done in Fig. 6E. Altogether, this requires F_p_(x) for neurons in the entire interactive group linear with a common slope, while this puts no limit on F_p_(x) for neurons in the noninteractive group, as shown in Eq. 7.

### Encoding failure and saturation

Another source of error is the amplitude of the input. In Fig. 2C, the amplitude was tuned such that the memory was graded across all neurons. However, if the amplitude goes smaller, with the overall shape unchanged, the low-firing neurons can have no memory, indicating *encoding failure*. An example is shown in Fig. S2A. If the amplitude goes smaller further, the input pattern may produce no memory, a total encoding failure. To avoid this, we require the input to activate at least one dendrite (another requirement is to sustain an activated dendrite for memory). The dendrite most easily activated is the one that projects from neuron 0 back to itself, having the largest pre- and postsynaptic firing rates I(0). We thus require:

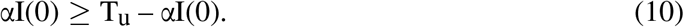

On the other hand, if the amplitude used in Fig. 2C goes larger, the memory starts to saturate. The saturation can happen in two ways. One is *out-degree saturation*, where high-firing neurons activate all dendrites they project to, across all neurons. This causes all neurons to have some activated dendrites, leading to a constant background firing (Fig. S2D). The other is in-degree saturation, where all dendrites in high-firing neurons are activated, saturating memory activity (Fig. S2G). See Appendix D for further details.

### Runaway and insensitivity

In the example shown in Fig. 3B, we mentioned that a positive feedback loop exists in the network dynamics. During the encoding period, this mechanism may destabilize the system, as the increase in firing rates can activate more dendrites, causing a recurrent increase in firing rates. This is the *runaway error*. Here, we provide a mechanistic analysis to describe the conditions under which this may occur. For the positive feedback loop during the memory period, see Appendix B.

To see how the encoding dynamics lead to the runaway error, we revisit Fig. 6B with an infinitesimal local increase δf(x_A_) at x_A_. This lowers 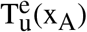 by αδf(x_A_), and increases the number of up-state dendrites by 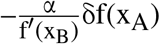. Therefore, the induced change in the summed dendritic output is:

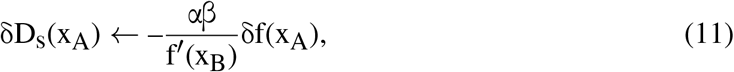

where δD_s_(x_A_) feeds back into δf(x_A_), leading to further changes in D_s_(x_A_). Above, we mentioned the in-degree error, which is due to the nonlocal dependency of the induced change on f^*′*^(x_B_). Differently, the runaway error happens when the gain factor, 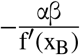, exceeds one, in which case the positive feedback causes the activity to blow up. To prevent the runaway error, we need:

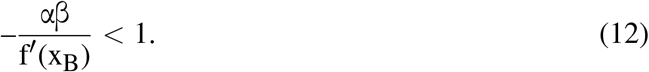

As 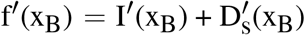, we have 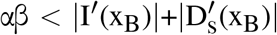. We get the sufficient condition to prevent the runaway error:

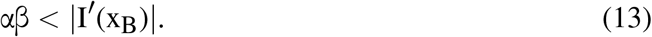

This condition should be applied to any x_A_, across all neurons. Therefore, the range of x_B_, which is geometrically determined by x_A_, includes all neurons in the interactive group. For the example in Fig. 6, the input satisfied this condition, and the memory had no runaway error. For the example shown in Fig. S3A, runaway error happened, but it did not completely destroy the graded memory.

On the other hand, we want D_s_(x) to be sensitive to a small increase. That is, δf(x_A_) leads to a nonzero change δD_s_(x_A_). This requires the gain factor to be nonzero, |f^*′*^(x_B_)|< ∞. This implies |I^*′*^(x_B_)|< ∞, such that there is no sudden drop in I(x_B_). Otherwise, the *insensitivity error* occurs, where part of the memory gives a flat firing rate, insensitive to graded input changes, as in the example shown in Fig. S3C. See Appendix E for more details about runaway and insensitivity errors.

### Memory performance with strong inhibition

In Fig. 6, we have examined network performance under an extra input increase during the encoding period. In parallel, here we examine network performance under an extra inhibition during the memory period and show how two functional groups of neurons differ in their responses to it. From a physiological perspective, such an inhibition might be performed by optogenetic silencing of a portion of a neuron population. From a cognitive perspective, people may need to temporarily silence one working memory representation to focus on a different task, and research shows that they can reactivate the temporarily silenced memory later [28].

Fig. 7 shows an example of how a maintained memory in our model reacts to an extra inhibition. After the memory was maintained, we applied a strong local inhibition to the soma for an extended period, after which we removed the inhibition. Fig. 7A shows the stable activities during this process for an inhibition targeting the interactive group of neurons. Upon the application of the inhibition, the original memory (dotted red line, reproduced from Fig. 2C) was silenced in the inhibited neurons (blue arrow), along with some activity reduction in other neurons (pink line). This reduction is explained by the corresponding dendritic states. Because the inhibited neurons had zero activity, their out-degrees fell to zero (Fig. 7B, blue arrow), and previously up-state dendrites (Fig. 2D) dropped to down-states. This deactivation led to a reduction in the memory activity beyond just the inhibited neurons, because other neurons lost some of their up-state dendrites that were originally sustained by the inhibited neurons. Notably, upon the removal of the inhibition, the memory in the inhibited neurons restabilized to a high level (Fig. 7A, red line). This is because up-state dendrites still existed in the inhibited neurons, supported by dendritic inputs from other neurons and not directly affected by the strong inhibition applied to the soma. Overall, similar to Fig. 3B where a graded memory was still maintained under large somatic noise, a graded memory can still be preserved after a temporary inhibition.

**Figure 7.**
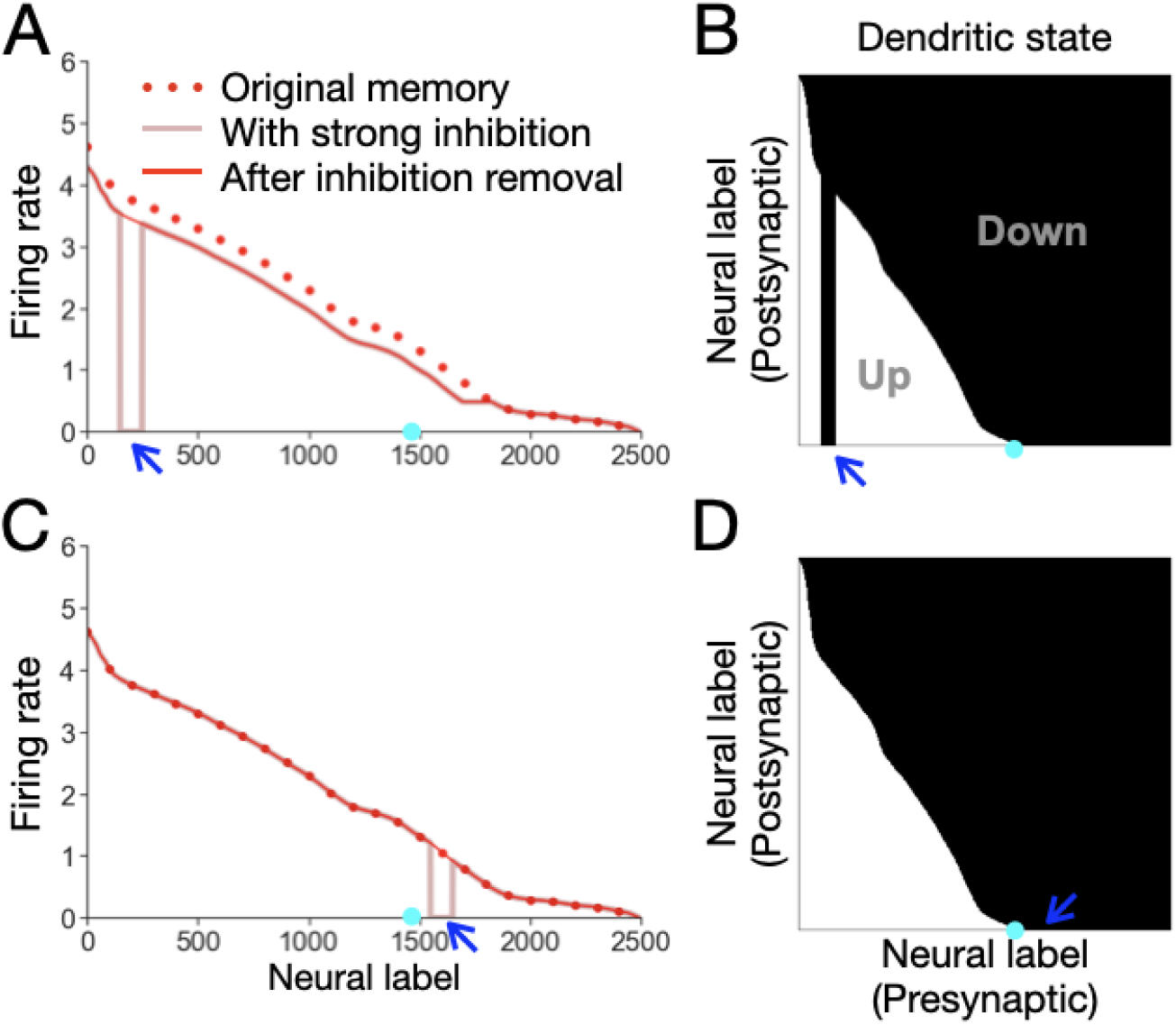
Memory performance when some neurons were briefly silenced. **A**, A strong local inhibition to the interactive group (left-hand side of the cyan dot). With the original memory in Fig. 2C (dotted red line), a strong local inhibition was applied to the interactive group of neurons. The inhibition killed the local activity (blue arrow) and reduced activity in other neurons (pink line). After its removal, the activity of inhibited neurons bounced back (red line). **B**, Dendritic states for the memory in *A* after inhibition removal. The inhibited neurons (blue arrow) having no out-degrees. The memory activity in *A* is proportional to the summed up-states along the vertical axis. **C, D**, Similar to *A, B*, but with the inhibition applied to the noninteractive group (right-hand side of the cyan dot). The memory was perfectly recovered after the inhibition removal.

In contrast, if the local inhibition was applied to neurons in the noninteractive group, where neurons having no out-degrees, the inhibition did not affect other neurons at all, and the inhibited neurons fully recovered (Fig. 7C, D).

More generally, memory reduction and recovery depend upon the in- and out-degrees of inhibited neurons. High-firing neurons have more out-degrees, and inhibiting them reduces broader activity across the network. Meanwhile, they also have higher in-degree and are more susceptible to dendritic deactivation because of the silencing of presynaptic neurons.

### The necessity of dendritic bistability for memory performance

Here, we check whether the bistable dendrite defined in Eqs. 2 and 3 is necessary for memorizing novel patterns. First, we set the bistable range to zero by redefining the down-threshold T_d_ in Eq. 2 as 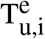. This makes each dendrite no longer follow a bistable relation, but rather a step function with a single threshold 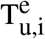. Therefore, instead of both Eqs. 4 and 5, the network dynamics can be described by Eq. 4 alone. In this case, the stable firing rate during the memory period 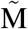 is given by

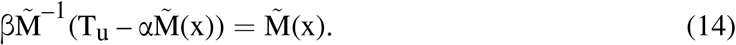

Note that this form is mathematically identical to Eq. 6 by choosing T_u_ = βx_0_. Therefore, its solution is also analogous to the solution of Eq. 6, which is Eq. 18: 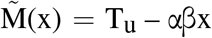, with x ranging from 0 to T_u_/β. However, this solution is unstable, as 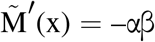 does not satisfy the stability requirement, Eq. 12.

On the other hand, if there is no somatic effect, with α = 0 in Eq. 3, all dendrites, in Eq. 2, across all neurons have the same 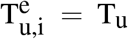 and T_d_ values. During the encoding period, for a given neuron, more dendrites can still be activated if the amplitude of the input pattern increases. However, across neurons, their 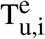 values no longer differ, such that they have the same number of activated dendrites and show the same memory activity. Fig. 8 shows an example of this case.

**Figure 8.**
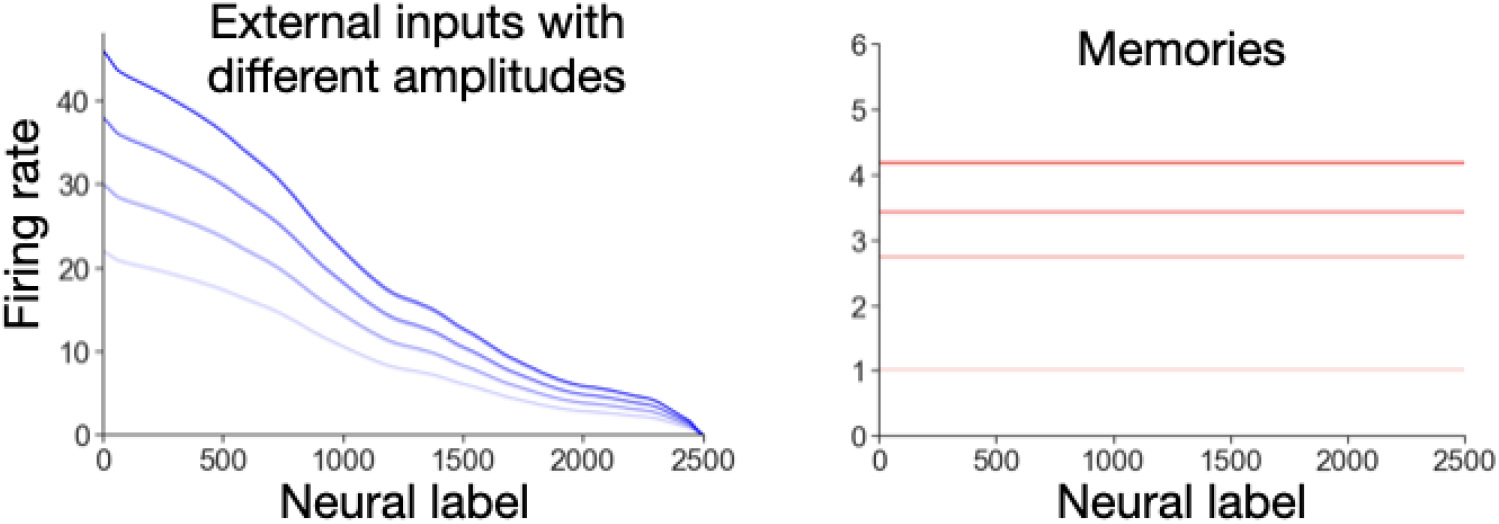
Memory performance in the absence of the somatic effect (α = 0). An input pattern gives a uniform memory. An input pattern with a larger amplitude gives a higher but uniform memory level, matched by the corresponding shading.

### A spiking network to maintain novel graded patterns

To demonstrate the biological plausibility of the rate-based model, we present a spiking network model to memorize novel graded patterns. The network consists of 40 randomly connected neurons, among which the probability of a connection is 50%. As shown in Fig. 9A, each connection projects to a separate dendrite, which is conductance-based with NMDA-receptor dynamics [19, 21]. On average, each neuron has 20 dendrites, whose voltages are summed, along with the corresponding external input, in the integrate-and-fire soma.

**Figure 9.**
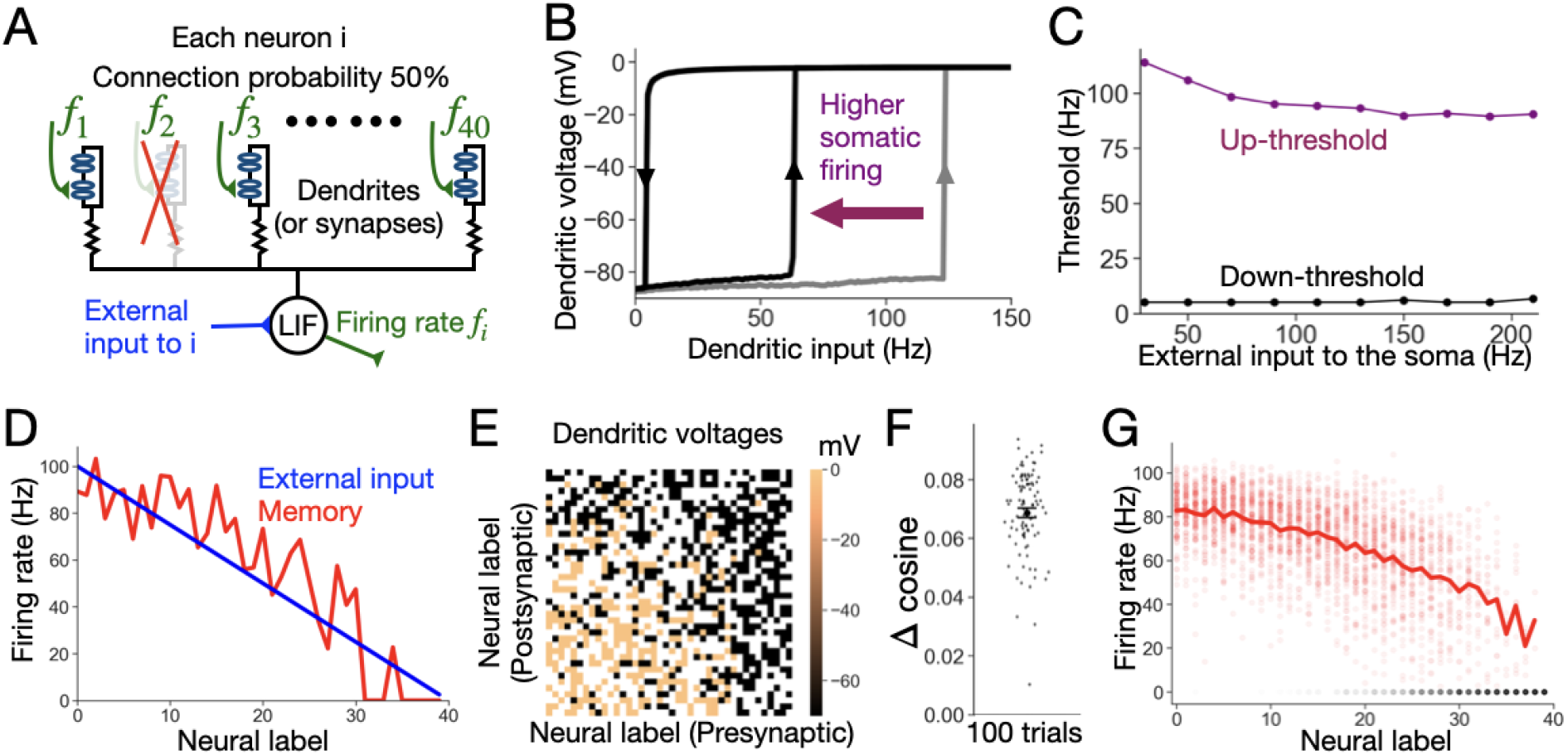
Spiking network model and its performance. **A**, Network structure. The network has 40 neurons. The connection probability between any two neurons is 50%. Each connection goes into a separate conductance-based dendrite (or, equivalently, synapse). The integrate-and-fire soma receives external input and all dendritic voltages. **B**, Dendritic bistable voltage. A dendrite shows bistable voltage with changing dendrite input frequency. Higher somatic firing can lower its up-threshold (purple arrow). Dendritic voltage was averaged over 500 ms for a smooth plot. **C**, How external input affects thresholds. Threshold values in *B* were recorded as the external input increased. Up-threshold dropped (purple), while the down-threshold remained mostly unchanged (black). **D**, Performance for memorizing a novel graded pattern. A linear input (blue) was delivered. This resulted in a noisy memory (red) with a similar graded pattern. **E**, Memory in dendritic voltages. The heatmap shows voltages of all dendrites of the network neurons, during the memory period in *D*. Each white pixel indicates the absence of a connection. **F**, Memory performance for 100 trials with the condition in *D*. The y-axis shows the amount of cosine similarity above the baseline value, which was calculated from a uniform memory, as in Fig. 3D. Each time, the network had a different initialization. The error bar represents the standard error of the mean. **G**, Explicit memory record of the 100 trials. The red dots represent non-zero memory activities for each neuron across trials. The red line represents the average of all non-zero memory activities, which shows a decreasing trend. The highly overlapping gray dots represent no memory activity. See Appendix F for more details.

The core dendritic mechanism in the rate model (Fig. 1C) was reproduced in this spiking network. Fig. 9B shows that a randomly chosen dendrite exhibits bistable voltage. In addition, increasing the somatic firing of its attached neuron will lower its up-threshold (purple arrow). Fig. 9C shows the average up-threshold reduction as the external input to the soma increased. This reduction can be explained physiologically. Initially, with a small dendritic voltage, the voltage-dependent conductance of NMDA receptors is low (due to the blockage of Mg^2+^). Therefore, a large dendritic input is required to activate the dendrite. However, bAPs, from somatic firing, can provide the depolarization to the dendrite to remove the Mg^2+^ blockage from the NMDA receptors, making their conductance larger and effectively lowering the required dendritic input for activation.

The down-threshold of each dendrite behaves differently. Fig. 9B,C indicates that the down-threshold is insensitive to the somatic firing rate. Physiologically, if the dendritic voltage is already depolarized, the conductance of NMDA receptors remains high (unblocked from Mg^2+^). Therefore, the additional depolarization from the somatic firing, via bAPs, has a relatively insignificant effect on the dendrite. In addition, the down-threshold is typically low as the simulations have shown. This is because the current contribution of NMDA receptors is easily saturated, which means that lowering dendritic input does not reduce the contribution much, keeping the dendrite still depolarized. This low down-threshold can be beneficial as, once activated, dendrites can easily be maintained even with a small dendritic input.

This spiking network can memorize novel graded patterns. Fig. 9D depicts the performance under an example linear input, with the memory maintaining graded values despite some noise. Fig. 9E shows the corresponding dendritic voltages during the memory period, which are higher for dendrites whose pre- and postsynaptic firing rates are high, showing the associative nature of each dendrite as in the rate-based model. Fig. 9F shows the memory performance across 100 trials, each with a different initialization. Each memory had a cosine similarity above the baseline. However, a subtlety exists due to the imperfection of this metric: a cosine similarity above the baseline does not necessarily indicate a graded memory. If the memory is only binary, with a lower memorization probability for neurons receiving low input rates, this also gives a cosine similarity higher than the baseline. Therefore, to further demonstrate that this network can store graded memory values, we recorded individual non-zero memory activities in Fig. 9F, as red dots. The average (red line) exhibits a decreasing trend, showing graded memory values. Additionally, neurons receiving lower input rates are more likely to have no memory activity, as indicated by overlapping gray dots. See Appendix F for more details.

## Discussion

In this paper, we present a network for the storage of novel graded patterns in working memory. In its simplest form, the network has uniform connectivity. Each neuron contains multiple bistable dendrites. The activation of each dendrite is associative, determined by pre- and postsynaptic firing rates. The number of dendrites in the up-state for a given neuron can vary, allowing the neuron’s firing rate to vary in a graded manner. We demonstrated the robustness of the network to various perturbations without fine-tuning, including somatic noise, connectivity perturbation, two-input interference, or a strong inhibition. This robustness was further demonstrated in a spiking network implementation of this model. We analyzed the dynamics of the network, showed the explicit condition for perfect memories, and identified various errors analytically such as in-degree error, out-degree error, encoding failure, encoding saturation, runaway error, and insensitivity. During this process, we further characterized how two functional neuron groups could emerge, with the interactive group having more impact to other neurons and thus requiring to be well constrained and the noninteractive group having no impact to other neurons.

Mechanistically, maintaining a novel graded pattern relies critically on local saturation in bistable dendrites. In previous models [14, 15] with fast Hebbian plasticity, upon receiving external input, the overall activity can easily saturate due to mutual excitation (a positive feedback loop). In our model, saturation happens locally in each dendrite, where increasing dendritic input no longer increases dendritic output. This local saturation prevents the positive feedback loop that could lead to global saturation of overall activity, allowing the system to still remain sensitive to different inputs. A similar concept is discussed in [29].

In addition, the somatic effect critically controls the ability to memorize novel graded patterns, as shown in Fig. 8. In other words, if the strength of the somatic effect can be actively controlled by some other mechanisms, the network can flexibly switch between a state capable of holding arbitrary patterns and a state that cannot. In the latter case, neurons may more flexibly interact for information processing, or neural activity is dominated by existing attractors for working memory. Consistent with this, it has been suggested that the strength of bAPs, as a main candidate for the somatic effect, may be controlled by dendritic-targeting inhibition [30, 31].

### Biological plausibility and alternative implementations

Dendritic bistability is essential to the present model. One way to achieve this bistability, as shown in the spiking implementation above, is to take advantage of the unique dynamics of NMDA receptors, as in previous works [19, 21]. The effect of the somatic voltage on the dendritic thresholds is a second essential component of the present model. One major candidate for this somatic effect is bAPs [32, 33, 34], as shown above. However, this is not the only possibility, especially when action potentials are missing, or the back-propagation is weak [35]. Other mechanisms may similarly provide the required postsynaptically dependent depolarization. For example, this can arise from subthreshold voltage in the soma or another input current roughly colocalized within the target dendrite [35]. Similar ideas have been tested in [36] for long-term potentiation. Conversely, if bAPs are very strong, dendrites may be highly coupled with the soma [37] with a potential worry that the soma and dendrites will act as a single unit instead of separate units. However, a previous model [38] shows that dendrites can still maintain high independence from the soma.

The spiking implementation of our model included an average of only 20 bistable dendrites per neuron, a realistic but small estimate which gives noisy performance. There are two potential ways to greatly increase the number of bistable units such that the performance is closer to the case in the continuum limit. One way is to use bistable synapses instead of bistable dendrites. Experiments [39, 40] have suggested that each synapse can function as a separate computational unit. A second way is to use bistable neurons in place of bistable dendrites. The dendrites of each neuron in the current model would map to a cluster of neurons, which would bidirectionally connect to another neuron, mapped by the soma in the current model. This implementation would yield a small-network level realization of the presented intracellular structure, where cellular bistability [41, 42] is required instead of dendritic bistability. In all of these cases, the key is to have a set of bistable units, which provide stability, and to combine them so that a graded value can be produced.

### Multiselectivity and two interpretations of the performance

The multiselectivity shown in network dynamics is a characteristic property of the model and makes neural firing rates correlated. It is rooted in the associative nature of the dendritic state, which is determined by both pre- and postsynaptic firing rates. During the encoding period, these associative dendrites may cause distortion in the formed memory and show an anticorrelation between susceptibility and influence among neurons as mentioned in Fig. 6E. However, the association can also be beneficial, as it helps the recovery of memory after temporary inhibition of a subset of neurons. Similar to synaptic plasticity rules, which maintain information in synapses, the memory maintains information in dendritic states, which may survive somatic inhibition.

All descriptions above by default consider the network as a single system. However, moti-vated by very different behavior in the interactive and noninteractive groups of neurons, especially for the case of perfect memory, we propose a second, dual-system interpretation. Suppose that the input consists of two separate sources simultaneously. One is an auxiliary internal input from a certain brain region, providing fixed high intensities to the interactive group. Another is an arbitrary external input, providing relatively low intensities to the noninteractive group. For example, the internal input may represent a general control or attentional signal of the current task setting, and the external input can represent a more explicit and flexible task-relevant sensory input. The sensory input can be memorized only if the internal input is active, providing neurons receiving the sensory input with large dendritic inputs during the encoding period. The final memory maintains activities caused by both the auxiliary input and the external input.

Adopting this dual-system interpretation can reduce much of the memory distortion. As it targets the noninteractive group, the arbitrary external input does not induce any out-degree error in other neurons. The induced memory for each neuron in this group is affected by the same slope of high firing rates as Fig. 6B shows, which is fixed internally. If the internal input is further trained to follow a linear decay in Eq. 7 (Fig. 5B), an arbitrary external input can be memorized perfectly. The potential price paid for this interpretation is that training could be required to provide a fixed internal input.

### Another way to memorize graded patterns, by parametric working memory models

There may exist another approach to storing novel patterns in working memory, in which each element of the input is stored in a separate simple circuit that is capable of storing a continuous scalar value (without any connections among these simple circuits) [5, 26, 27]. In other words, the same simple memory circuit is independently replicated for each input element. There have been many models for memorizing a single scalar value in structure attractors, also known as parametric working memory [5,26,27], with both cellular [41,42,43,44] and network mechanisms [4, 6, 7, 45, 46].

It is questionable whether this replication approach can store novel information in working memory. Such parametric working memory models require structured attractors for scalar memorization. It is possible that such attractors are gene-encoded without prior training, especially for working memory related to evolutionarily important intuitions. However, it is not likely that all information, such as language, concepts, or objects, has corresponding attractors in the brain predefined by genes. In other words, this approach likely needs attractor formation and is not suitable for novel working memory.

There are also distinctions in memory performance between our model and the replication approach. Our model shows multiselectivity, which is more consistent with recordings during working memory tasks [47, 48] in regions with dense mutual connections. In contrast, the replication approach maintains each input entry separately, exhibiting single selectivity with no connections or interactions among replicas.

In summary, we propose a network model based on dendritic bistability that is capable of representing novel graded patterns in working memory.

## ACKNOWLEDGMENTS

This work was supported by the Air Force Office of Scientific Research (AFOSR) under award number FA9550-22-1-0532.

## Appendix

### A: Simulation details of the rate-based network model

Details of the rate-based network implementation are included here. For the default settings in Eq. 1 and Eq. 2, parameters are N = 2500, T_u_ = 20, T_d_ = 1, β = 0.0032, α = 0.7, τ = 50 ms and w = 1. Firing rates are non-negative and equal to the sum of the dendritic contribution and the external input. To memorize a novel input pattern, the network started with all dendrites in the down-state. An input was applied during the encoding period for 1000 ms, and it was turned off during the following memory period for another 1000 ms. Equilibrium was reached in each period.

The default input pattern, which first appeared in Fig. 2, was generated based on a cloud image N083 (CCSN·v2/Cu/Cu-N083.jpg) in [49]. The original image was compressed to a resolution of 50×50, with grayscale. The original intensity took an integer value from 0 to 255, with degeneracy. Therefore, we added a small value to each intensity to get rid of this degeneracy. The input pattern had the minimal intensity subtracted and an amplitude of 15.33. Eventually, this resulted in discrete input values I_i_ with i from 1 to N = 2500.

In Fig. 2B, we calculated the cosine similarity between external input patterns with different amplitudes and the corresponding memories. The input amplitude ranged from 10 to 40 in steps of 0.5. The baseline value was calculated based on a uniform memory. We also checked an alternative baseline using the cosine similarity between the input and the shuffled memory with bootstrapping. This gave a lower baseline value, with no qualitative changes to our results.

Simulation details for Fig. 3A-D are included here. In Fig. 3A,B, we implemented independent Gaussian noise throughout the memory period. The noise has mean zero and σ one (Fig. 3A) or 5 (Fig. 3B), applied with a time step of 1 ms. The dark red line was obtained by averaging the memory activity over the last 500 ms during the 1500 ms memory period.

The noise-free memory in Fig. 2C was reproduced as the dotted red line for comparison. In Fig. 3C, we implemented four variations to the original connectivity. First, weight values were drawn from a Gaussian distribution with mean 1 and σ 0.5, bounded below by zero. Second, the connection probability p was changed to 10%. The total number of connected dendrites was reduced, and the unconnected dendrites were never activated, effectively being deleted from the network dynamics. To balance this reduction, β was normalized correspondingly: β ← β/p. Third, with the original network, we added a local attractor with a radius of 15, centered at the middle of the two-dimensional pixel space. The weight value between any two neurons within the attractor equals two. Fourth, each neuron was projected to a random dendrite of the targeted neuron, such that a dendrite could receive zero to multiple dendritic inputs from network neurons. For all panels in Fig. 3C, we used the same scale bar for ease of direct comparison. However, the actual firing rates of some neurons could exceed the maximum scale used. This nuance does not change qualitative results. In Fig. 3D, we recorded memory performance for 100 input patterns (N001 to N100). For each input pattern, we generated performance similar to that in Fig. 2B for a range of input amplitudes, from 10 to 32 in steps of 0.5. The optimal input amplitude, which gave the maximum cosine similarity, was used for that input pattern across all conditions. For conditions with noisy connectivity, a different connectivity initialization was used for each data point.

Simulation details for Fig. 3E,F are included here. In Fig. 3E, input 1 (N026) was delivered in the first encoding period, followed by the first memory period. Then, input 2 (N084) was delivered, followed by the second memory period. The right or left 50×20 region of input 1 or 2 was set to zero. The uniform memory used to compute the baseline was also limited to the left or right region. In Fig. 3F, we ran 100 pairs of such inputs. Input 1 ranged from N001 to N100, and input 2 was different from input 1, randomly selected between N001 and N100. In addition, the same sets of input 1 and 2 were run by themselves alone, without interference from the other for comparison. Input amplitudes were taken from Fig. 3D, with an extra 5 added (to reduce the impact of input activity drop due to setting the left or right region to zero). There were still about 3 choices of input 1 or 2 that could not be memorized by themselves alone (zero memory and negative Δ cosine similarities). In addition, there were about 5 simulations in the two-input interference case that had negative Δ cosine similarities.

Simulation details for the rest of the main text figures are included here. In Fig. 5, the external input used for a perfect memory has a linear part I_p_(x) = 14.4 – 0.00576x, for x between 0 and A = 1500. For the arbitrary part, the simulated input values were taken from the lowest 1000 intensity values of a different image (N004). This portion was rescaled so that its maximum value matched the minimal value of the linear part in neuron 1500. In Fig. 6, an extra input was given to neurons 1700 to 1750 (Fig. 6A) or 1300 to 1350 (Fig. 6C) during the encoding period with a magnitude of 1 added to the default I(x). In Fig. 7, a signal with an intensity of −20 (which was large enough to completely silence local firing rates) was given to neurons 150 to 250 (Fig. 7A) or 1550 to 1650 (Fig. 7C) at the end of the memory period. It lasted for 1000 ms, and another 1000 ms was simulated following its removal. In Fig. 8, we set α = 0, and the input amplitudes used are I(0) = 22, 30, 38 or 46.

Simulation details for the supplementary figures are included here. In Fig. S1A, T_d_ = 2. In Fig. S2A, D, and G, the input amplitudes are I(0) = 13, 17, and 19, respectively. In Fig. S3, the derivative was estimated using F(x) values from four nearby neurons: F^*′*^(x) ≈ [–F(x + 2Δx) + 8F(x + Δx) – 8F(x – Δx) + F(x – 2Δx)]/(12Δx), with a step size of Δx = 5. This numerical differentiation for discrete values gives a better approximation of the true derivative in the continuum limit. In Fig. S3A, we chose the same image as used in Fig. 5, with an input amplitude I(0) = 14.17. In Fig. S3C, the default input pattern had an amplitude I(0) = 15.8, and all intensities after neuron 1200 were decreased to 80% of their original values, so that discreteness was created in the input pattern at neuron 1200.

Simulations were performed using the forward Euler method with a step size of dt = 1 ms in Python [Version: 3.9.16].

### B. Forgetting during the memory period

**Figure S1:**
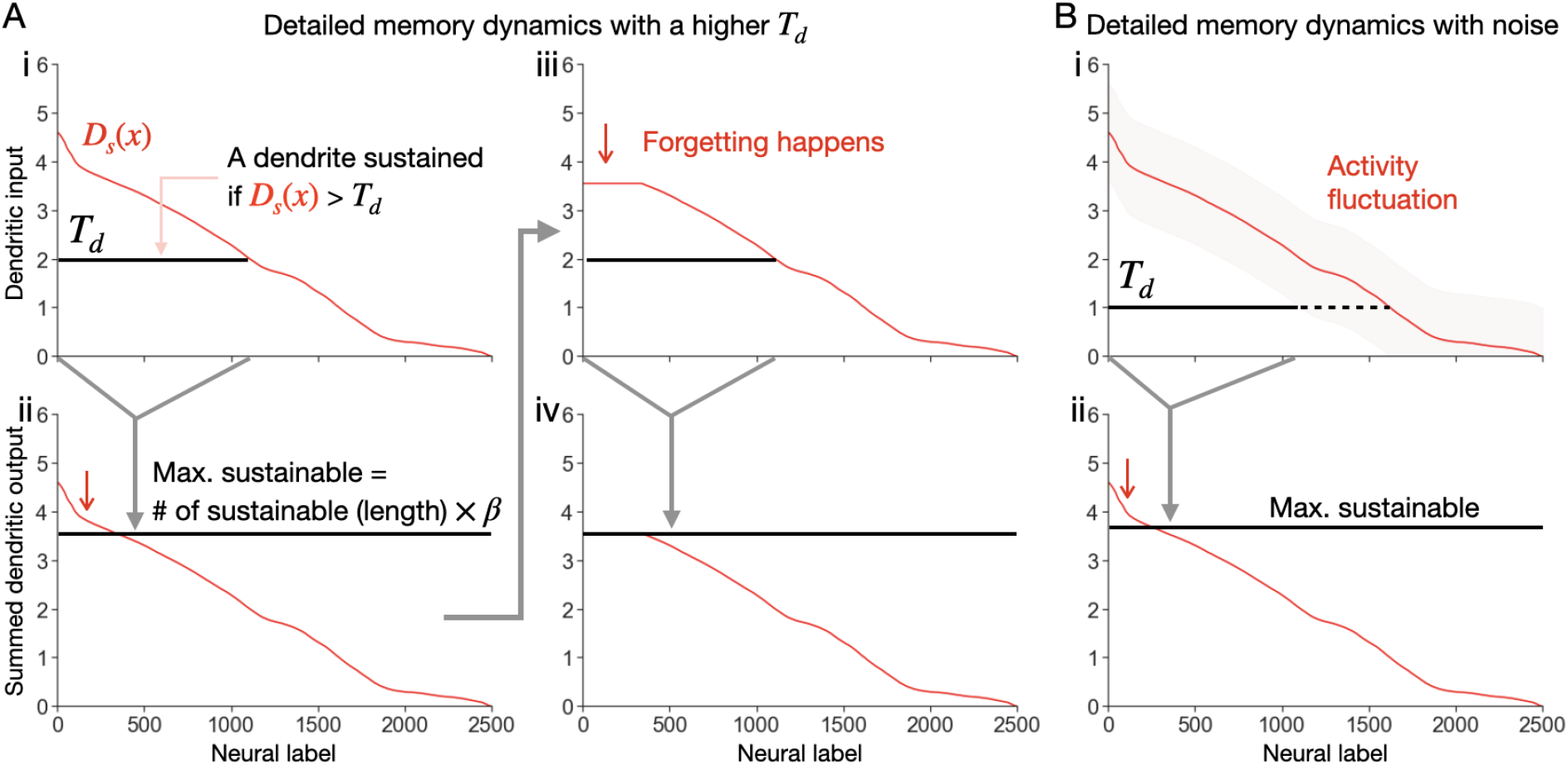
Detailed memory dynamics with forgetting. The same dynamics as Fig. 4 are shown, but with a higher T_d_ or noise. In these conditions, Fig. 4H, I, are replaced by the plots here. **A**, Case with a higher T_d_. i, The black line shows the portion of dendrites with dendritic inputs above T_d_. ii, The maximum sustainable activity (black line) is lower than in Fig. 4I. Activated dendrites in high-firing neurons drop to the down-state and reduce activity (red arrow). iii, The updated D_s_(x) is used to determine sustainable dendrites iteratively. The length of the black line is unchanged. iv, The maximally sustainable activity is unchanged. D_s_(x) is now sustainable, with M(x) = D_s_(x). **B**, Case with noise during the memory period. i, Noise makes activity fluctuate (shaded). The solid line shows the portion of dendrites always receiving inputs larger than T_d_ despite the noise. Dendrites in the dashed line region can go below T_d_ due to noise fluctuations. ii, High-firing neurons drop memory activity (red arrow) as in the noisy memory example shown in Fig. 3B.

In Fig. 4G-I, we described the memory dynamics when no forgetting happens. Here, we extend the dynamics to the case of forgetting. That is, once-activated dendrites during the encoding period fall back to the down-state. In this case, some dendrites are more susceptible than others. Based on the example in Fig. 4, we used a higher T_d_. The dynamics are different, with Fig. 4H,I being replaced by Fig. S1A. Compared with the previous case in Fig. 4H, in Fig. S1Ai, a higher T_d_ makes the number of sustainable dendrites smaller (black line). Some up-state dendrites in high-firing neurons are not sustainable (Fig. S1Aii, red arrow), and forgetting happens there (Fig. S1Aiii). Importantly, this decrease in activity may not lead to further recurrent decrease in memory. This is because the number of sustainable dendrites is determined by the intersection of T_d_ and low firing rates, which may be unchanged even when forgetting happens, as shown in Fig. S1Ai, Aiii. That is, forgetting may clip high-firing neurons, but not low-firing neurons, as the stabilized memory in Fig. S1Aiv shows.

For memory degradation under large noise (Fig. 3B), high-firing neurons are clipped similarly. But instead of having a higher T_d_ as in Fig. S1A, here activity can be temporarily lowered due to noise, as indicated by the shaded fluctuation region in Fig. S1Bi. This makes the number of sustainable dendrites smaller, and thus leads to forgetting. Note that in general, it is not easy to flip a down-state dendrite to the up-state, which requires noise to be comparable to the external input strength during the encoding period.

### C. Solving for a perfect memory

We derive conditions for a perfect memory here. We first simplify Eq. 6 to

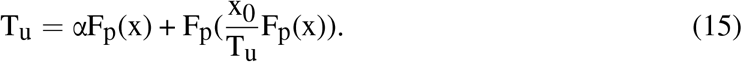

Recall that F_p_(x) is a monotonic function that decays to 0, F_p_(N) = 0. Substituting the relation into Eq. 15 gives: F_p_(0) = T_u_.

On the other hand, taking the derivative of Eq. 15 gives:

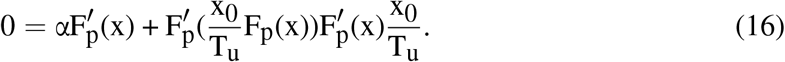

For a nontrivial solution, 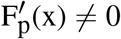, we have:

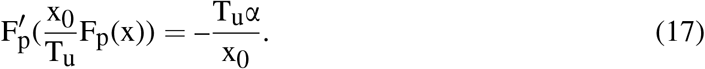

This equation gives a constant slope 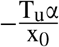 for F_p_(x) with a range determined by 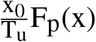. Note that F_p_(x) can take values from F_p_(N) = 0 to F_p_(0) = T_u_. This means that 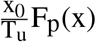 takes values from 0 to x_0_. In other words, F_p_(x) is linear with a slope 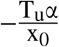 for x from 0 to x_0_.

Altogether, we have the solution for Eq. 6:

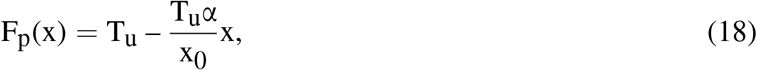

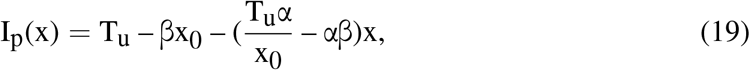

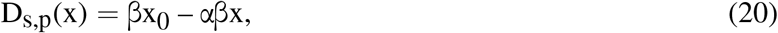

where x ranges from 0 to x_0_. We require 1 ≥ α > 0 such that F_p_(x) is not negative for x = x_0_.

In addition, I_p_(x) needs to be large enough, such that it can activate some dendrites during the encoding period. This requires the input amplitude, I_p_(0) = T_u_ – βx_0_, to satisfy:

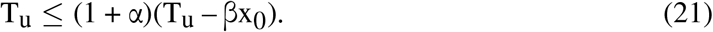

Otherwise, total encoding failure happens, as mentioned above with Eq. 10.

On the other hand, to prevent any forgetting, we need 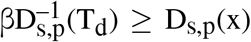, as mentioned above in Fig. 4I. As D_s,p_(x) has a maximal value of βx_0_, we have 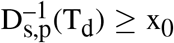. With Eq. 18, we get:

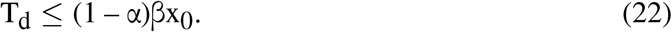

Recall that 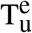 has a lower bound, T_d_. Is this lower bound reached in the perfect memory case? The lowest 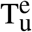 exists in the highest-firing neuron x = 0 with F_p_(0) = T_u_, having a value of T_u_ – αT_u_. We then have T_u_ – αT_u_ > (1 – α)βx_0_ ≥ T_d_, which shows no saturation in 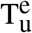. Note that for the first inequality, we use the relation T_u_ > βx_0_, which is derived from Eq. 21, 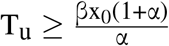.

Altogether, we get the condition for a perfect memory in the main text.

### D. Further details about how input amplitudes lead to encoding failure and saturation

**Figure S2:**
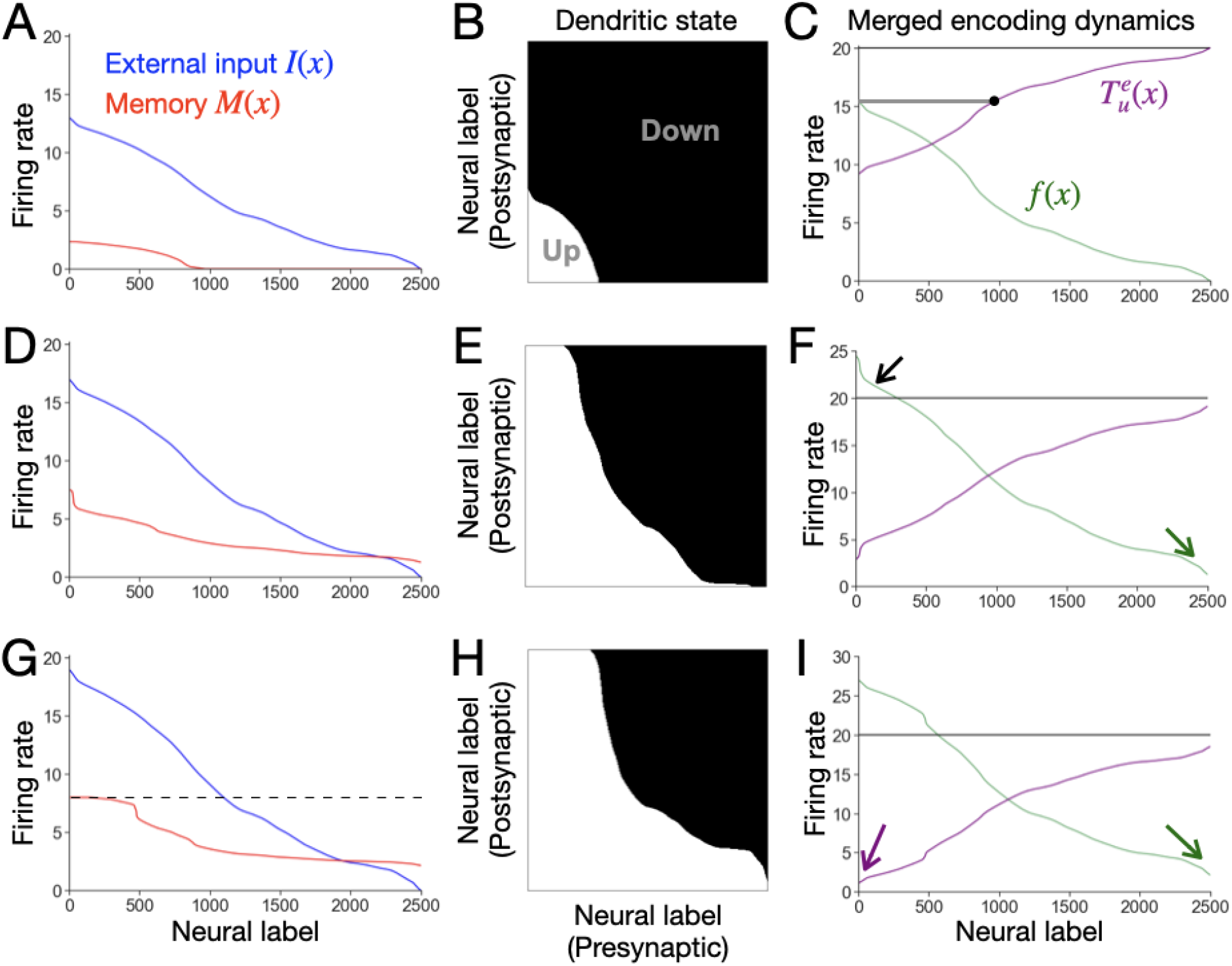
Encoding failure or saturation. **A-C**, Memory with encoding failure. **A**, An example. The input used has a smaller amplitude, compared to the one used in Fig. 2C. This resulted in a memory in which neurons received low input intensities remained silent. **B**, Dendritic states of the memory. **C**, Merged encoding dynamics. For low-firing neurons (right-hand side of the dot), their 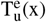 values are too high to allow dendritic activation in these neurons, resulting in zero memory. **D-F**, Memory with out-degree saturation. **D**, An example. The input pattern used has a larger amplitude, compared to the one used in Fig. 2C. This resulted in a constant background memory activity. **E**, Dendritic states of the memory. **F**, Merged encoding dynamics. The high-firing neurons exceeding T_u_ (black arrow) activate all dendrites they project to, regardless of the somatic effect. Therefore, every neuron has dendrites activated by these high-firing neurons, leading to constant background activity (green arrow). **G-I**, Memory with in-degree saturation. **G**, An example. The input pattern used has an even larger amplitude, compared to the one used in *D*. This resulted in further memory activity saturation (dashed horizontal line). **H**, Dendritic states of the memory. **I**, Merged encoding dynamics. Even the smallest firing rate (green arrow) can exceed the 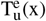 values of high-firing neurons (purple arrow). Therefore, all dendrites in these neurons are activated.

Eq. 10 shows the necessary condition to avoid total encoding failure. But what is the condition to avoid any encoding failure? That is, for a neuron receiving a nonzero input value, we want a nonzero memory value. This requires the neuron receiving an infinitesimal input (i.e., neuron N, with 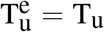) to have at least one dendrite activated. The dendrite most easily activated in this neuron is the one that receives dendritic input from neuron 0, with a maximum value of f(0). Therefore, we need:

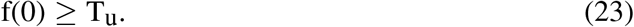

The input in Fig. 2C was tuned to make F(0) = T_u_ such that no encoding failure happened, where F(0) is the stabilized, maximal value of f(0) during the encoding period. If this inequality is not satisfied, some neurons receiving small input values (low-firing neurons) will have no activated dendrites, leading to no memory. An example is shown in Fig. S2A. During the encoding period (Fig. S2C), there is a dot determined by the horizontal line from the maximum firing rate f(0). For neurons on the right-hand side of the dot, their 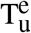 values are higher than the maximum firing rate f(0), such that none of their dendrites are activated, giving no memory.

On the other hand, Fig. S2D shows an example of out-degree saturation. The high firing rates exceed T_u_ during the encoding period (Fig. S2F, black arrow), thereby activating all dendrites that they project to (Fig. S2E). These up-state dendrites, distributed across all neurons, provide a constant background activity (Fig. S2F, green arrow). Out-degree saturation is benign, as a constant background still keeps graded memory values.

In contrast, in-degree saturation can be more problematic. It can happen in high-firing neurons in two ways: 1, their 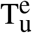 values hit the same minimum T_d_; 2, their 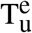 values are lower than all dendritic inputs received. In either case, these neurons have the same number of activated dendrites, leading to memory saturation. An example of the latter case is shown in Fig. S2G-I, where 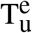 values for the high-firing neurons (Fig. S2I, purple arrow) exceed the lowest firing rate (green arrow).

To avoid encoding failure and saturation, the parameter range needs to be 0 < α ≤ 1. Avoiding encoding failure requires f(0) ≥ T_u_ as mentioned above. Based on this, if α > 1 instead, the 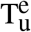 of the highest-firing neuron hits the minimum value 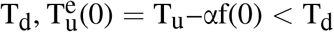, which leads to in-degree saturation.

### E. Further details about how input slope determines runaway and insensitivity errors

**Figure S3:**
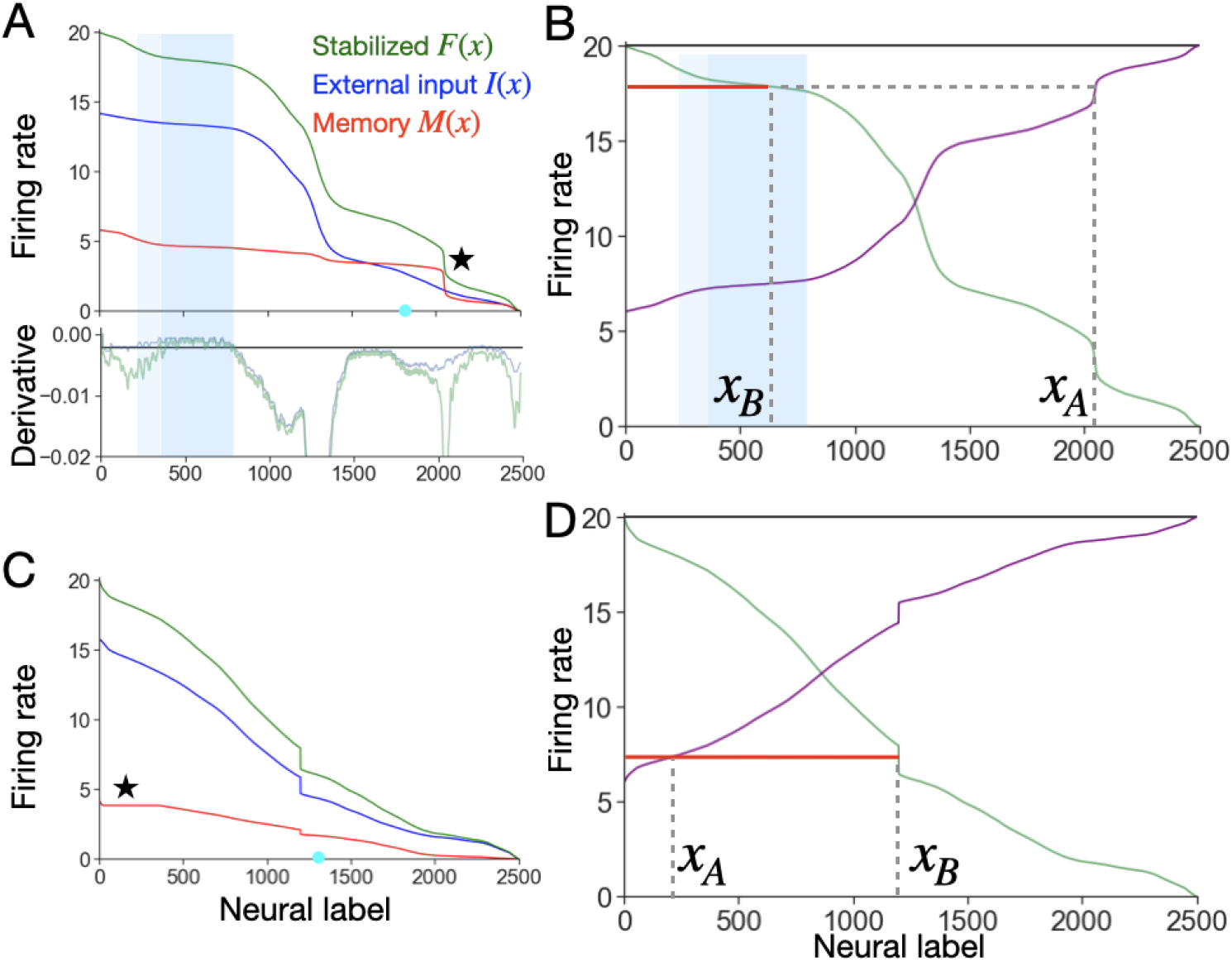
Runaway and insensitivity errors. **A, B**, Runaway error. **A** Top: Basic performance. The green curve shows the stabilized firing rate F(x) during the encoding period. The cyan dot separates the interactive group (left) and the noninteractive group (right). Bottom: Derivatives of F(x) (green) and I(x) (blue). A horizontal line indicates the value of –αβ. The lighter blue region has derivatives of I(x) above –αβ. The darker blue region has derivatives of F(x) above –αβ. A local discrete jump in the memory exists (star), even though I(x) has no jump. **B**, Merged encoding dynamics. **C, D**, Insensitivity error. **C**, Basic performance. The input has a discrete jump in the middle, making the memory discrete at the same position. In addition, the memory shows a flat region (star). **D**, Merged encoding dynamics. See details in the text.

Fig. S3A shows an example of the runaway error where I(x) or f(x) failed to satisfy Eq. 12 or Eq. 13. f(x) was chosen to be the stabilized firing rate F(x) during the encoding period. Fig. S3A, bottom, shows the slopes of I(x) and F(x), with the horizontal line showing –αβ. The violation of the sufficient condition, Eq. 13, may not lead to the runaway error as long as Eq. 12 is still satisfied (light blue region). However, the violation of Eq. 12 (dark blue region) gives the runaway error with a local activity blowup (starred). As the encoding dynamics show (Fig. S2B), the dendritic inputs from neurons in the dark blue region to neuron x_A_ are close in value. That is, a small increase δf(x_A_) can lead to the activation of all these dendrites through a positive feedback loop, resulting in a discontinuous jump in D_s_(x_A_).

Fig. S3C shows an example of the insensitivity error. The input I(x) used has a discrete drop. As the encoding dynamics show (Fig. S3D), a slightly larger activity in neuron x_A_ does not change the in-degree of x_A_, the length of the red line, because of the discrete jump in x_B_. This leads to a flat memory in Fig. S3C (starred).

For runaway and insensitivity errors, we only consider how a local increase applied to neuron x_A_ directly causes x_A_ to gain more up-state dendrites. Technically, the local increase may cause more activity in other neurons first, which acts back on x_A_ for more activation. This indirect feedback pathway is ignored because an increase in x_A_ maximally activates one dendrite in other neurons. In the continuum limit, the contribution of a single dendrite is negligible.

### F. Simulation details about the spiking network

The spiking network structure was described in the Results. Each neuron consists of an integrate-and-fire soma with n conductance-based dendrites, adapted from previous models [19, 21]. The following equations show the dynamics for each neuron j. For clarity, we omit the neural label j.

The somatic voltage V_s_ follows dynamics:

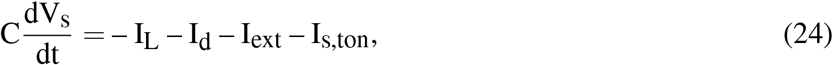

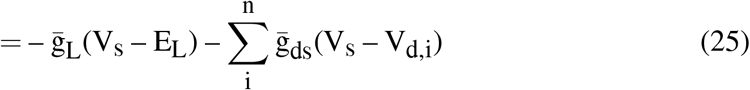

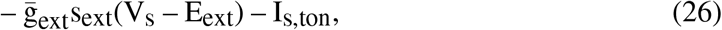

where I_L_ is the somatic leak current, I_d_ is the total current from all dendrites, I_ext_ is the current from an external spike train, and I_s,ton_ is the tonic background input. The membrane capacitance is C = 10 nF/mm^2^. The maximal conductance for the leaky term is 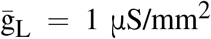, for the coupling from dendrites to the soma is 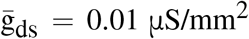, and for the external input is 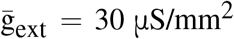. The equilibrium voltages are E_L_ = –80 mV and E_ext_ = 0 mV. I_s,ton_ is independent Gaussian noise with mean –34 nA/mm^2^ and σ 1 nA/mm^2^, applied every 0.25 s. n is taken to be the actual number of dendrites for each neuron, with mean 20, given a connection probability of 50%. The synaptic activation s_ext_ is specified below. Once V_s_ exceeds the firing threshold –50 mV, a spike is triggered, and the somatic voltage is set to 30 mV for 3 ms before being reset to –55 mV for another 3 ms.

The dendritic voltage V_d,i_ for a dendrite i follows dynamics:

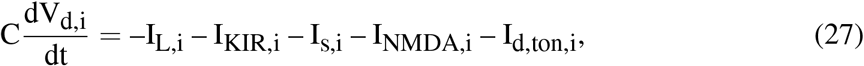

where I_L,i_ is the dendritic leaky current, I_KIR,i_ is the inward-rectifying potassium current, I_s,i_ is the current from the soma, I_NMDA,i_ is the current driven by dendritic input through NMDA receptors, and I_d,ton,i_ is independent Gaussian noise with mean zero and σ 1 nA/mm^2^, applied every 0.25 s. The terms on the right-hand side follow:

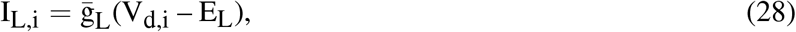

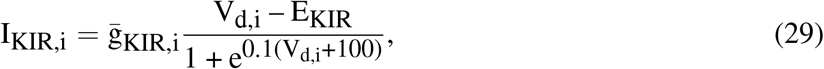

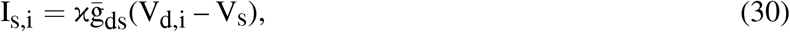

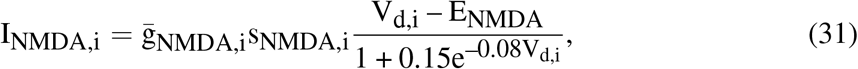

where the equilibrium voltages are E_NMDA_ = 0 mV, E_KIR_ = –90 mV, and the area ratio of the soma to a dendrite is κ = 50. The maximal conductance 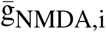 is a Gaussian random variable with mean 54 μS/mm^2^ and σ 1 μS/mm^2^. Similarly, 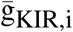 with mean 62 μS/mm^2^ and σ 1 μS/mm^2^.

The synaptic activation s_α_ follows:

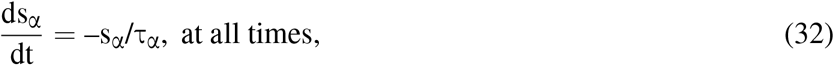

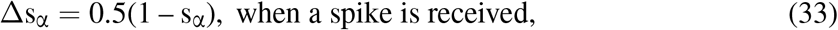

where s_α_ represents synaptic activation for the external input (s_ext_) or dendritic input to dendrite i (s_NMDA,i_), with a decay time τ_ext_ = 2 ms or τ_NMDA,i_ = 100 ms, respectively. s_ext_ receives an external spike train, different for each neuron, and s_NMDA,i_ receives the dendritic input from other neurons i, if a connection exists.

With the above dynamics, each dendrite can show a bistable curve in voltage. To show that this dendritic bistability is rooted in each dendrite, instead of a network-level effect, we considered an isolated neuron. That is, for Fig. 9B, C, we chose a neuron with n = 20 dendrites and removed all its recurrent connections. These dendritic inputs were instead provided by a fixed-period spike train with varying frequency. This qualitatively mimicked dendritic inputs from other neurons, which have a relatively fixed period due to integrate-and-fire somas. To trace out a bistable curve, we increased and then decreased this frequency while recording the changing dendritic voltage. Specifically, the input frequency to the dendrites started from 0 Hz and increased by 1 Hz every 1500 ms until it reached 150 Hz. Afterwards, it decreased by 1 Hz every 1500 ms until it went back to 0 Hz. The recorded dendritic voltage was averaged over the last 500 ms before each frequency change. The gray curve in Fig. 9B was generated with an additional somatic input, a Poisson spike train with a frequency of 30 Hz. The black curve was generated with a frequency of 210 Hz instead. For Fig. 9C, we ran 10 trials for each somatic input and recorded the averaged threshold values across all 20 dendrites over all trials. Threshold values were estimated from the largest slopes. The somatic input ranged from 30 Hz to 210 Hz in steps of 20 Hz.

In Fig. 9D-G, we simulated the homogeneous network with 40 neurons. Voltages were stabilized at about –80 mV initially. The input to each soma was a Poisson spike train with a frequency linearly decreasing from 100 to 0 Hz as shown in Fig. 9D. The encoding and memory periods lasted for 1000 ms. In Fig. 9D,E, the recorded dendritic voltage and firing rate were averaged over the last 500 ms. In Fig. 9F, we ran 100 trials, each with a different initialization. The cosine similarity was calculated between the fixed linear input in Fig. 9D and the resulting memory. We further subtracted the baseline cosine similarity, a constant of 0.87 (between the fixed linear input and a uniform memory). There was one trial with the performance below the baseline. In Fig. 9G, we recorded the explicit firing rates over these 100 trials.

Simulations were performed using the forward Euler method with a step size of dt = 0.25 ms in Python [Version: 3.9.16].

